# Resource competition shapes CRISPR-mediated gene activation

**DOI:** 10.1101/2024.07.03.601429

**Authors:** Krishna Aravind Manoj, Domitilla Del Vecchio

**Affiliations:** Department of Mechanical Engineering, Massachusetts Institute of Technology; Department of Biological Engineering, Massachusetts Institute of Technology

## Abstract

CRISPR-mediated gene activation (CRISPRa) allows concurrent transcriptional activation of many genes and has found widespread use in genome-wide screening, bioproduction, and therapeutics. Scaffold RNAs (scRNAs) recruit dCas9 and an activator protein (RBP-AD) to the target gene for activation with high sequence specificity. Here, we show that, despite this specificity, orthogonal scRNAs interfere with each other because they compete for dCas9 and RBP-AD. Specifically, we demonstrate that the expression of an scRNA that binds to these resources results in repression of genes targeted by different scRNAs. Intriguingly, we also discover that transcriptional gene regulation by an scRNA is biphasic, wherein increased level of the scRNA leads to gene repression instead of activation. These effects are significant even when dCas9 and RBP-AD are expressed at the maximum level tolerable by the cell. Our results demonstrate that CRISPRa systems are not as modular as previously thought and establish predictive modeling tools to assess the emergent behavior of multi-module CRISPRa networks.

---

CRISPR has revolutionized the field of genetic engineering and has transformed our ability to edit and regulate genomic loci in several organisms ^1^. In a CRISPR Cas9 system, the Cas9 protein binds to a guide RNA that helps locate the target gene with high specificity. In its native context ^2, 3^, this acts as an adaptive immune system response where the Cas9 protein cleaves the DNA of viruses infecting the bacteria ^4^. DNA modification through the CRISPR Cas9 system has then moved from its original function of protecting bacteria to genomic manipulations ^5– 9^ and to control gene expression using catalytically inactive dead Cas9 (dCas9) ^10^. In CRISPR dCas9 systems, guide RNAs recruit the dCas9 protein to specific genes based on predictable Watson-Crick base pairing. Specifically, during CRISPR-mediated gene interference (CRISPRi), the guide RNA recruits dCas9 to physically block RNAP binding to the target promoter site, thereby repressing transcription ^10^. In contrast, CRISPR-mediated gene activation (CRISPRa) also uses an activator protein that recruits RNAP to the promoter site to enhance transcription. In this case, the guide RNA includes an additional hairpin loop for the recruitment of the activator protein and is called scaffold RNA (scRNA)^11, 12^. CRISPRi and CRISPRa, therefore, enable simultaneous and orthogonal regulation of multiple genes, which has been used for genome-wide screening ^13– 15^, bioproduction ^16– 18^, and therapeutic applications ^19, 20^.

Although CRISPR/dCas9 systems are designed in a modular fashion and can in principle allow concurrent and independent regulation of multiple targets, this is not the case in practice for CRISPRi ^21– 23^. In fact, dCas9 is shared by multiple guide RNAs, and this causes competition for this resource, wherein increased expression of one guide RNA causes a decrease in the dCas9 pool available to other guide RNAs. Since these effects cannot be mitigated by increasing the level of dCas9 due to its toxicity ^24^, a dCas9 regulator was designed to neutralize competition, restoring the modularity in CRISPRi systems ^22^. Although mathematical models of competition have appeared for CRISPRa systems ^25, 26^, it remains to be determined whether competition effects appear in practice.

To address this question, we created CRISPRa systems containing either one or multiple CRISPRa modules and assessed the effects of competition for dCas9 and RBP-AD on both on-target and off-target responses. We further evaluated the extent to which dCas9 and RBP-AD are each bottlenecks in CRISPRa, and demonstrated that increasing the levels of these resources does not mitigate the observed competition effects. We finally investigated the use of competition as a design parameter to tune the response of a CRISPRa module.

## Results

### On-target CRISPRa response is biphasic

To characterize the on-target response of a CRISPRa system, we created a CRISPRa module where we varied the concentration of the input scRNA and monitored gene expression output through a fluorescent reporter (Fig. 1a,b). CRISPRa is enabled by a short target-specific scaffold RNA (scRNA) that recruits dCas9 and RNA-binding protein with activation domain (RBP-AD) to activate a target gene ^11, 12, 27^. The scRNA consists of three distinct regions: a 20 bp complementary region for target gene binding, a hairpin region for dCas9 protein binding, and an MS2 hairpin for RBP-AD binding. The RBP-AD unit consists of the MS2 coat protein (MCP) fused with SoxS (R93A/S101A) that binds to the MS2 hairpin of the scRNA ^12^. The sequence of the CRISPRa target site is encoded in the 20 bp complementary region on the scRNA (Fig. 1a).

**Figure 1.**
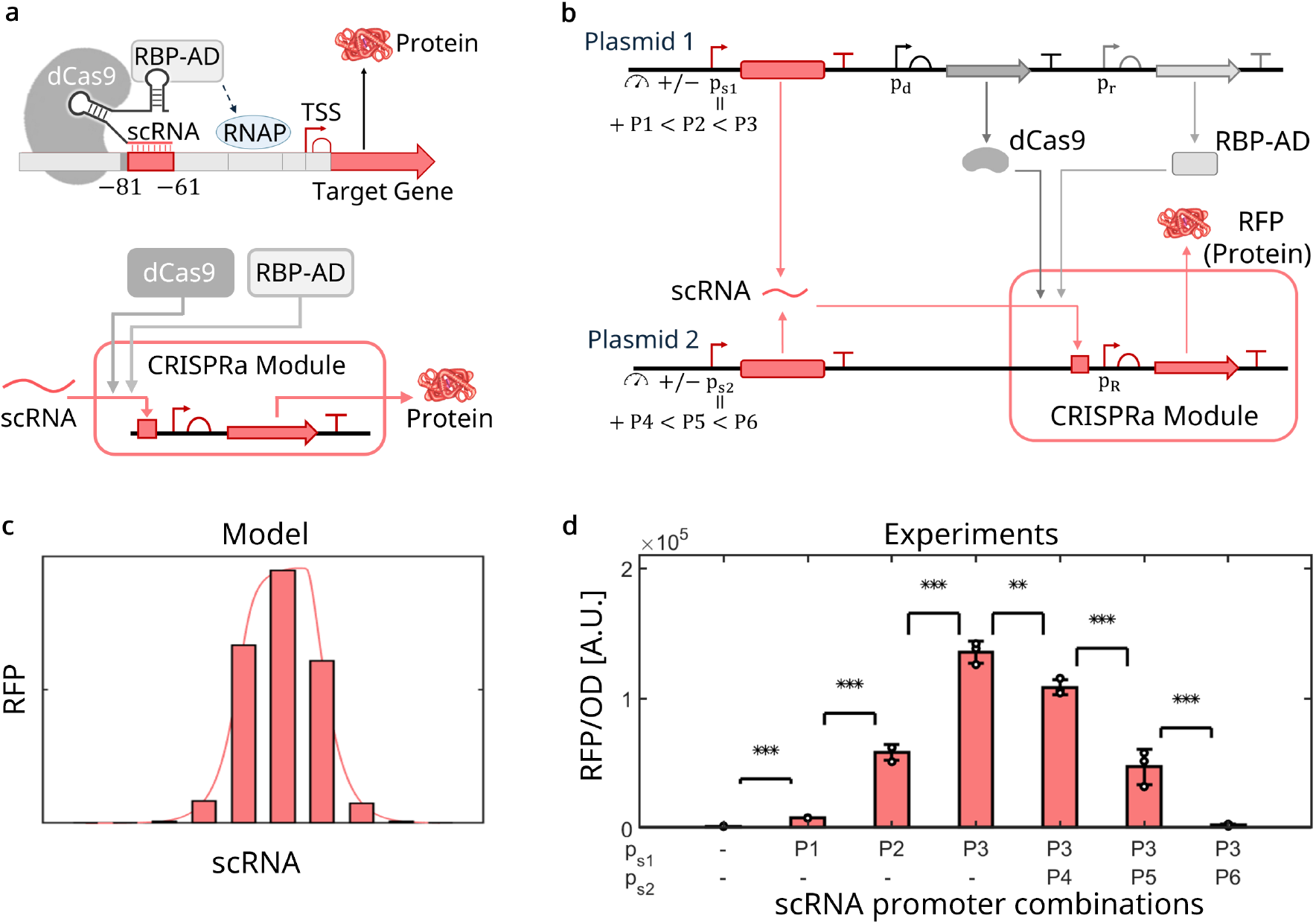
Biphasic on-target response of a CRISPRa module. (a) Genetic diagram showing the interactions between the scRNA and the two resources dCas9 and RBP-AD. (Top) The CRISPR complex consisting of dCas9, RBP-AD, and scRNA binds upstream of the transcription start site (TSS) of the target allowing activation. RBP-AD recruits RNAP to the minimal promoter, activating the expression of the target gene. (Bottom) Block diagram representation of a CRISPRa module with scRNA as the input and protein as the output. (b) Genetic construct of a CRISPRa module, where the scRNA is expressed with variable promoter strength from two sites and recruits dCas9 and RBP-AD to the promoter expressing RFP (see SI Note **1** for details and SI Fig. **1** for plasmid maps). (c) Mathematical model of the input/output response of the CRISPRa system in panel (b) (see SI Note **3** for the model). (d) Experimental data showing RFP protein level as the scRNA level is increased. The scRNA is increased by increasing the strength of its promoters p_s1_ and p_s2_ as indicated (see SI Fig. **2** for promoter strength quantification and SI Fig. **3** for the temporal data).

In our implementation, the scRNA binds upstream of the promoter (p_R_) expressing the RFP output protein (Fig. 1b). In order to vary the scRNA level over a wide range, we expressed it from two different DNA locations under the control of promoters whose strength varies according to a published library of promoters ^28^(SI Fig. **2**), while dCas9 and RBP-AD are constitutively expressed. We created a mathematical model in which we explicitly accounted for a finite total amount of dCas9 and RBP-AD proteins in the system (SI Note **3**). This model shows a surprising biphasic input/output response of the CRISPRa module (Fig. 1c); that is, while at low scRNA levels an increase in scRNA expression leads to increased output protein level, this is not the case at high scRNA levels. At high scRNA levels, increased expression of the scRNA leads to target repression and ultimately quenches the output completely. The experimental data confirms this finding and shows that at the highest expression levels of the scRNA, the output RFP protein is completely repressed (Fig. 1d).

**Figure 2.**
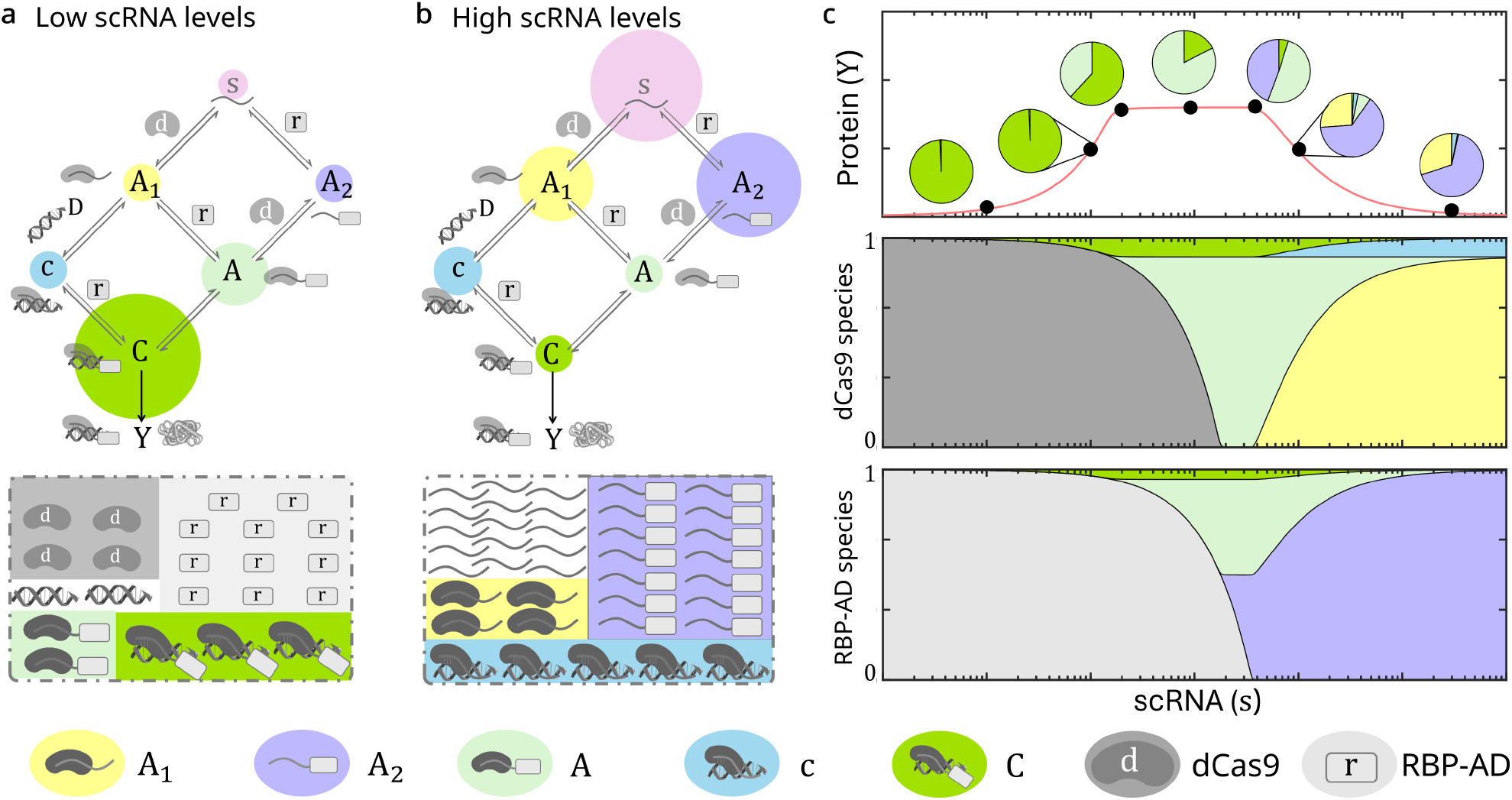
A double-diamond reaction diagram explains the observed biphasic CRISPRa response. (a) Schematic representation of the fractional concentrations of each species in the double-diamond reaction diagram of a single CRISPRa module for low scRNA levels showing the availability of free resources and the dominance of complexes with both resources. Similar representation of the fractional concentrations for high scRNA levels showing the abundance of scRNA and the depletion of complexes with both resources. (c) Numerical simulations showing the evolution of the relative abundance of individual species during the biphasic response of a single CRISPRa module. (Top) Biphasic response of the output protein *Y* with respect to its input scRNA *s* (red curve), the pie charts show the fractional concentration of each species in the reaction diagram in panels (a) and (b) for a given value of the input (black dot). Pictorial representations of fractional concentrations of species involving (Middle) dCas9 and (Bottom) RBP-AD, respectively, with respect to the input scRNA (*s*).

We thus interrogated our mathematical model to shed light on a potential molecular mechanism explaining this biphasic input/output response (Fig. 2). The model indicates that the biphasic response emerges because the scRNA is required to bind both dCas9 and RBP-AD before it can lead to the transcription of the target gene. Therefore, when the dCas9 and RBP-AD resources are in excess compared to the scRNA, the system is predominantly composed of the transcriptionally active complex (C in Fig. 2a), leading to an increase in the production of the protein as the scRNA is increased. By contrast, when the scRNA is in excess with respect to the levels of free resources, we have an abundance of complexes formed by the scRNA with either of the two resources (*A*_1_, *A*_2_ and *c* in Fig. 2b) leading to a lack of complexes with both resources (*A* and *C* in Fig. 2b). This, in turn, quenches output expression. As we increase the scRNA, we observe three distinct behaviors of the output protein levels: increasing, plateauing, and decreasing (Fig. 2c). Specifically, the concentration of the transcriptionally active complex and therefore the output (*C* and *Y* in Fig. 2a, respectively) increases when scRNA level increases but only until the dCas9-RBP-AD-scRNA complex (*A* in Fig. 2a) has saturated the target DNA, leading to the plateauing of the curve. Further increase of the scRNA level leads to the sequestration of RBP-AD and dCas9 from the dCas9-RBP-AD-scRNA complex into scRNA-dCas9 and/or scRNA-RBP-AD complexes (*A*_1_ and *A*_2_ in Fig. 2b). This, in turn, leads to a decrease in the transcriptionally active complex RBP-AD-dCas9-scRNA-DNA (*C* in Fig. 2b), which decreases the output level. At very high scRNA levels, we observe the complete sequestration of resources into the intermediate complexes, *A*_1_ and *A*_2_ (Fig. 2c and SI Fig. 4). This causes the output protein level to drop, leading to the observed biphasic CRISPRa response.

### Independent CRISPRa modules are coupled through resource competition

We next investigated the role of dCas9 and RBP-AD resource sharing in the ability of two CRISPRa modules to function independently (Fig. 3a). To this end, we constructed a genetic system with two independent CRISPRa modules: one module (Module 1) takes scRNA_RFP_ as input and gives RFP as output, while the second module (Module 2) takes scRNA_GFP_ as input and gives GFP as output (Fig. 3b). The GFP scRNA (scRNA_GFP_) is expressed by a constitutive promoter with variable strength, whereas the RFP scRNA (scRNA_RFP_) is expressed by a DAPG-inducible promoter ^29^. Our mathematical model accounting for the two modules and the conservation of resources (SI Note 3) shows that as we increase the level of DAPG, RFP increases as expected while GFP decreases, demonstrating an unintended coupling between the two modules (Fig. 3c - top). Thus, we performed experiments in which we increased scRNA_RFP_ by increasing the inducer level DAPG and observed that as RFP is activated, the GFP level decreases by approximately 46%, in accordance with the model predictions (Fig. 3c - bottom). This confirms a coupling between the two modules, wherein expressing the input scRNA to Module 1 results in a decrease in the output of Module 2. We also note that this coupling persists for different levels of the input scRNA of Module 2 (see SI Fig. 9). We further observed that these effects are reciprocal since increasing the scRNA_GFP_, input to Module 2, results in an 85% decrease of RFP, output of Module 1 (Fig. 3d, model on top and experimental data on the bottom). These effects persist, although to a lower extent, when the DAPG level and hence the input scRNA to Module 1 is increased (see SI Fig. 10). This experiment conducted in the absence of the GFP coding region in Module 2 with the strong scRNA_GFP_ promoter shows no significant difference between the RFP signals, demonstrating that these effects are not arising from ribosome competition ^30^ (SI **Fig. 11**).

**Figure 3.**
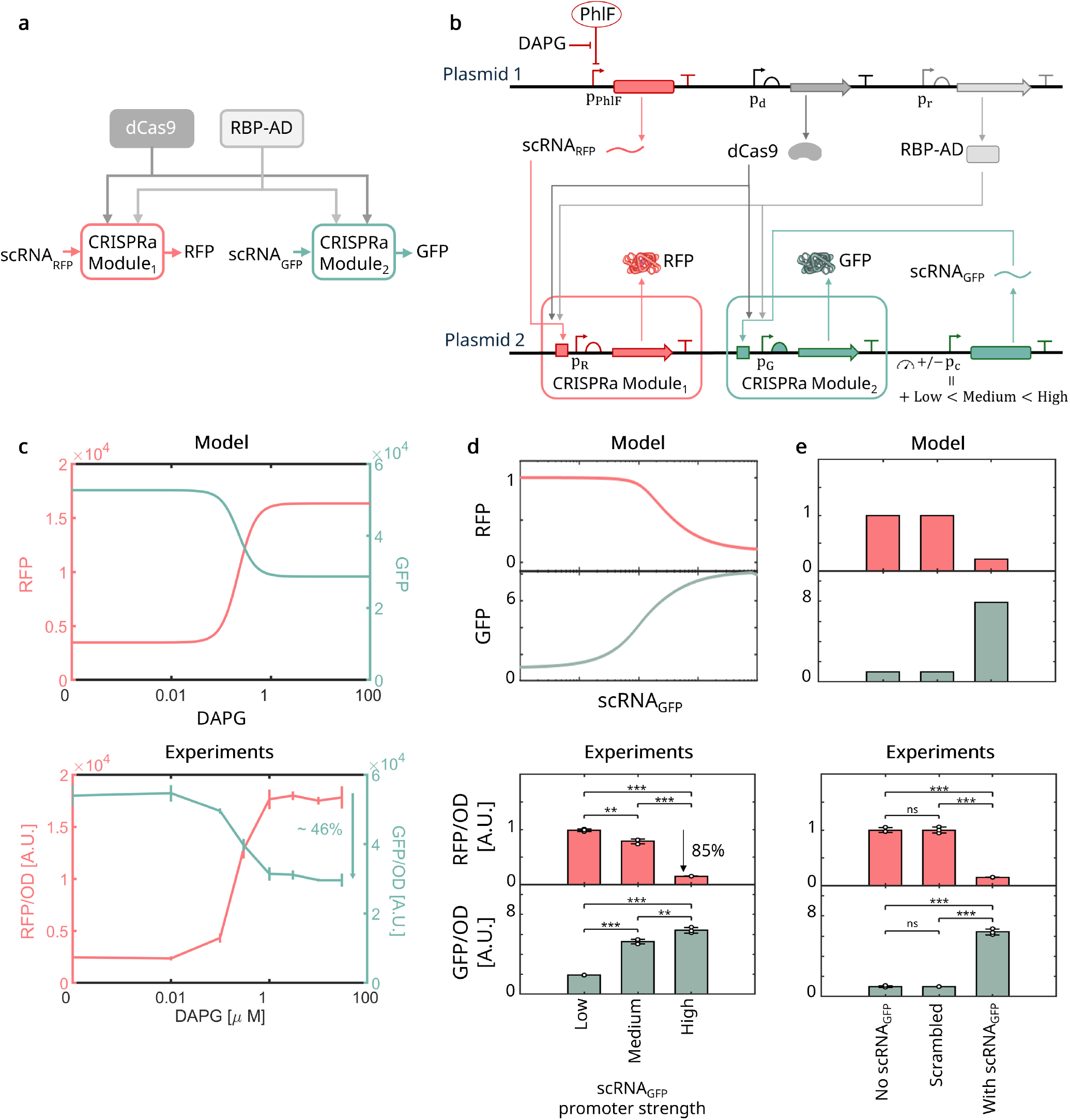
Coupling between two independent CRISPRa modules through dCas9 and RBP-AD resource competition. (a) Block diagram representation of two CRISPRa modules with their inputs (scRNA), output (proteins), and resources (dCas9 and RBP-AD). (b) Genetic construct with two CRISPRa modules. One takes scRNA_RFP_ as input and gives RFP as output; the other takes scRNA_GFP_ as input and gives GFP as output (plasmid maps and sequences are provided in SI Notes **1** and **4**). (c) On-target response of RFP and off-target response of GFP as scRNA_RFP_ level is increased via DAPG. (Top) Numerical simulations from the ODE model with two CRISPRa modules (see SI Note **3**). (Bottom) Experimental results with two CRISPRa modules using the genetic circuit in (b). (d) Fold change in RFP and GFP as the scRNA_GFP_ level is increased. (Top) Numerical simulations showing the response of RFP and GFP with continuous increase in scRNA_GFP_. (Bottom) Experimental data showing RFP and GFP for varying promoter strengths of scRNA_GFP_ as low, medium, and high (J114, LacUV5, and pTrc, respectively, SI Fig. **5**) for 0 DAPG. The charts are normalized using the protein levels obtained in the absence of scRNA_GFP_. (e) Fold change in RFP and GFP for three conditions of scRNA_GFP_: No scRNA_GFP_, scrambled scRNA_GFP_ which does not bind to dCas9 or RBP-AD and with perfect scRNA_GFP_. (Top) Numerical simulations from the ODE model for the three conditions (see SI Note **3**). (Bottom) Experimental results showing RFP and GFP for the three conditions with pTrc promoter for 0 DAPG (all scRNA sequences are provided in SI Note **4.2**). Time Series corresponding to the experimental data in (c)-(e) are in SI Figs. **6 - 8**.

To determine whether the observed coupling between the two modules is due to competition for dCas9 and RBP-AD, we designed two additional constructs where the scRNA_GFP_ is either absent or expressed by the strongest promoter but with a scrambled sequence, removing the binding with dCas9 and RBP-AD. We observed that the level of RFP is not affected by the presence of the scrambled scRNA_GFP_ and is the same as that obtained in the absence of the scRNA_GFP_ (Fig. 3e). This indicates that it is not the process of expressing the scRNA that causes the coupling between the two CRISPRa modules, but rather the coupling emerges from the binding of the scRNA to the RBP-AD and dCas9 resources.

### Engineering selective resource demand to identify dominant competition effects

We next investigated the relative contributions of the competition for dCas9 and RBP-AD on the observed coupling between the two CRISPRa modules. To this end, we designed the scRNA_GFP_ to selectively bind to either dCas9 or RBP-AD, thereby leading to sharing only one resource at a time between the two CRISPRa modules (Fig. 4a). The scRNA_GFP_ in Module 2, which we refer to as the competitor, consists of three distinct regions, the target gene (DNA), dCas9, and RBP-AD binding sites. We modified the scRNA_GFP_ to implement four different cases. In the first case (Case I), scRNA_GFP_ is a perfect competitor (the same as used in Fig. 3), which binds to both dCas9 and RBP-AD. In the second case (Case II), the dCas9 binding site on the scRNA_GFP_ is scrambled such that the scRNA only binds to RBP-AD but not to dCas9. In this case, the scRNA is also unable to bind with the target gene since dCas9 is required for binding to the DNA ^12^. In the third case (Case III), we scrambled the RBP-AD binding site on the scRNA_GFP_ such that the scRNA binds only dCas9 but not RBP-AD, thereby competing for the former but not for the latter. In the fourth case (Case IV), we scrambled both the dCas9 and RBP-AD binding sites of scRNA_GFP_, leading to no competition for resources between the two modules. The scRNA_GFP_ sequences for all four cases are in SI Note 4.2.

**Figure 4.**
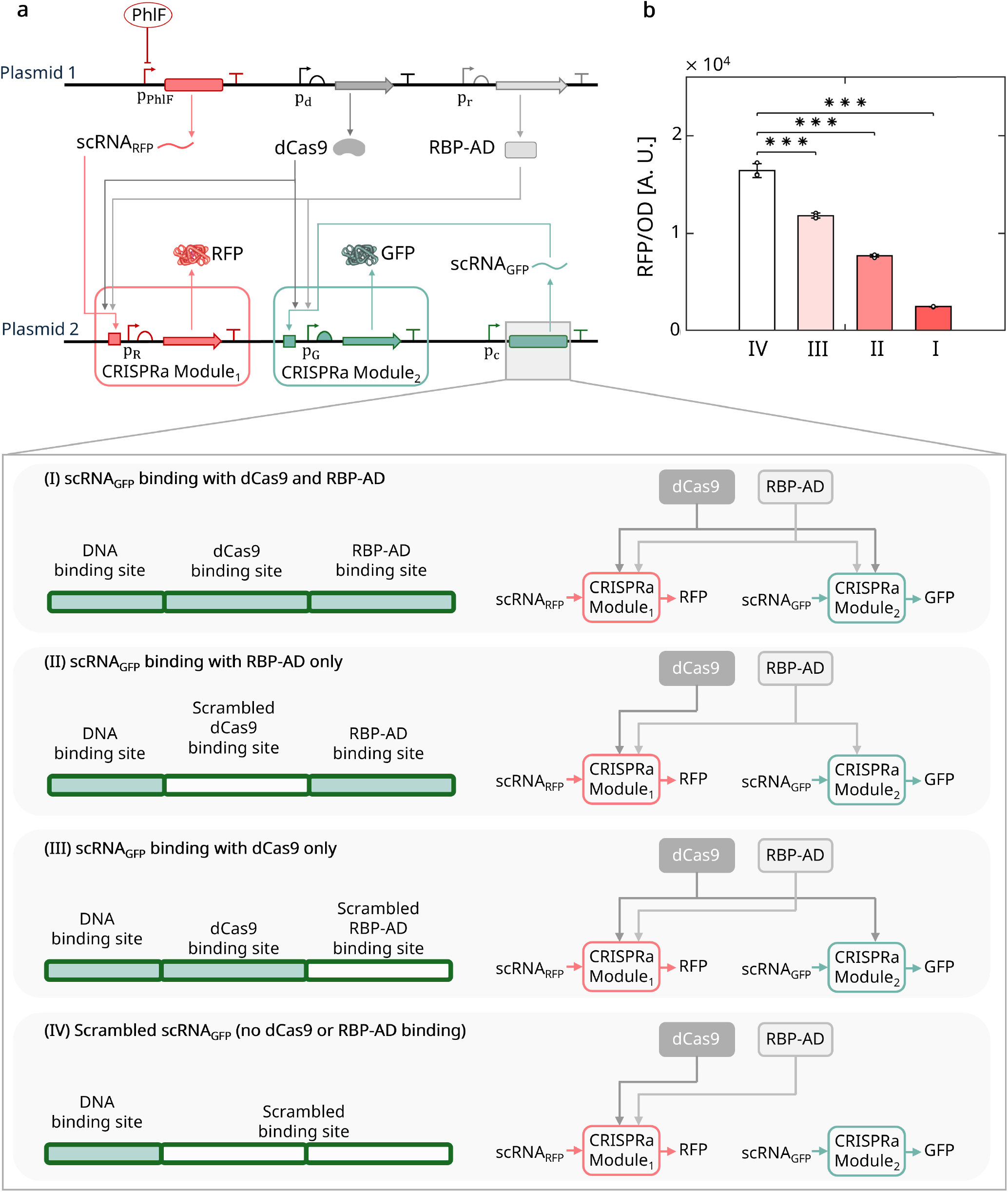
Competition effects arise from sharing both dCas9 and RBP-AD. (a) Genetic constructs of two CRISPRa modules with scRNA_RFP_ activating RFP and scRNA_GFP_ activating GFP. The scRNA-GFP is modified to selectively bind to dCas9 and/or RBP-AD: in (I) it binds with both dCas9 and RBP-AD, in (II) it binds only to RBP-AD, in (III) it binds only to dCas9, and in (IV) it binds to neither RBP-AD or dCas9 (all sequences are in SI Note 4.2). The block diagrams indicate how the resources are shared between the two CRISPRa modules for the different scRNA_GFP_ variants. (b) Experimental data showing the output of Module 1 when the scRNA_GFP_ is expressed in each of the four cases in panel (a) for 0 DAPG. Temporal data is in SI Fig. 12.

We observed that when the competitor scRNA binds to either dCas9 or RBP, but not to both, the RFP level still drops, although not to the same extent as observed when the scRNA binds to both resources (Fig. 4b). This indicates that the observed competition effects are not due to predominant competition for one or the other resource but for both of them. Furthermore, when both resources are shared between the two modules, the effects of competition are more significant compared to when only one of them is shared. These effects are observed independent of the level of the input scRNA_RFP_ to Module 1 (SI Fig. **13**) and are recapitulated by the model (SI Note **3.1** and SI Fig. **14**).

### Increasing resource level does not mitigate competition

We next investigated whether increasing the levels of resources is a viable solution to mitigate the effects of competition. Therefore, we examined the extent to which the biphasic shape of the on-target response of a CRISPRa module and the coupling between two CRISPRa modules persist as the dCas9 level is increased. To this end, we first created a genetic construct for a single CRISPRa module in which the dCas9 promoter strength is variable and we examined the response of the CRISPRa module’s output as dCas9 level is increased (Fig. 5a). Surprisingly, the model indicates that as the level of dCas9 is increased, the output RFP increases for low dCas9 levels but then decreases for high dCas9 levels (Fig. 5b-left). Experiments with varying promoters for dCas9 confirmed this finding and that the maximal expression level of RFP occurs for intermediate levels of dCas9 instead of being reached for maximal dCas9 expression (Fig. 5b-right). Hence, the output of the CRISPRa module does not monotonically increase with dCas9 level, leading to a biphasic response to dCas9. According to our model, this biphasic response is due to the limitation of scRNA and RBP-AD for high dCas9 levels (SI Fig. 17).

**Figure 5.**
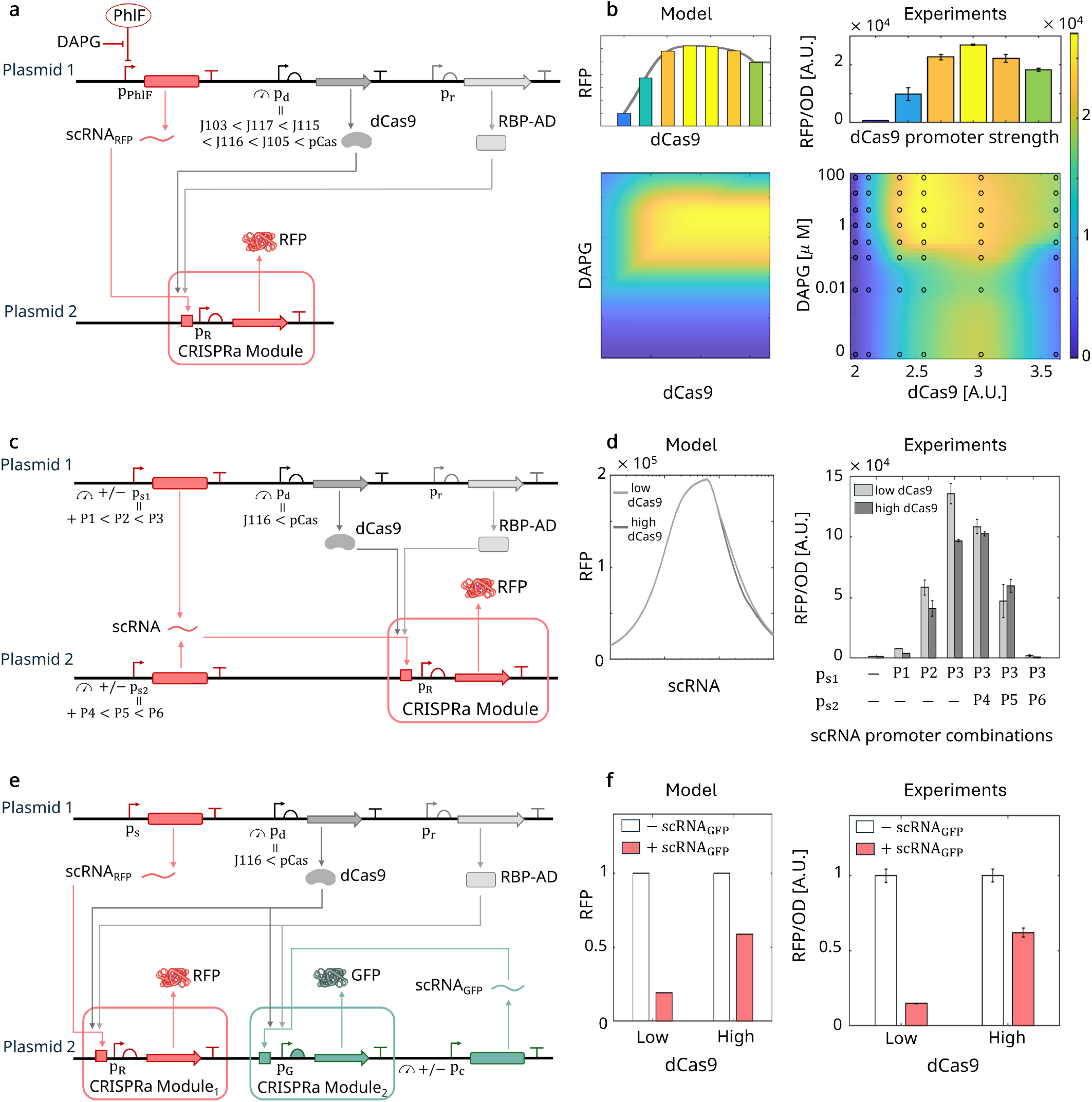
Increasing dCas9 level does not remove competition effects. (a) Genetic construct for a single CRISPRa module with scRNA_RFP_ activating RFP with varying promoter strengths for dCas9. The promoters were taken from the Church library ^28^ and are given by BBa_J23103 (J103), BBa_J23117 (J117), BBa_J23115 (J115), BBa_J23116 (J116), BBa_J23105 (J105) and pCas (see SI Note 4 for the sequences and SI Fig. 15 for strength characterization). (b) Effect of dCas9 level on RFP. Bar chart shows RFP with increasing dCas9 promoter strengths at 30*µ*M DAPG. Left: Mathematical model showing RFP for varying levels of dCas9 and scRNA_RFP_. Right: Experimental data showing the output RFP levels for different dCas9 promoters and DAPG levels. The circles are experimental measurements colored by the mean fluorescence values of three biological replicates (standard deviations in SI Fig. 16). The continuous color in the background is the interpolation of the experimental data. (c) Genetic construct with a CRISPRa module with input scRNA_RFP_ and output RFP for two different promoters expressing dCas9 (p_d_ given by J116 and pCas promoters, see SI Fig. 15). (d) Response of RFP to input scRNA_RFP_ for different dCas9 levels. Left: Mathematical model of the input/output response of the CRISPRa module shown in (c). Right: Experimental data. (e) Genetic construct with two CRISPRa modules with scRNA_RFP_ activating RFP and scRNA_GFP_ activating GFP for p_d_ given by J116 (low) and pCas (high) promoters (see SI Fig. 15). (f) Bar chart representation of the fold-change in RFP in the absence (−scRNA_GFP_) or presence (+ scRNA_GFP_) of the competitor scRNA for low and high dCas9 levels. Left: Numerical simulation results showing RFP with and without the competitor scRNA for different levels of dCas9. Right: Experimental data showing the output of Module 1 with and without the input for Module 2 for different promoter strengths for dCas9.

We thus next investigated whether the biphasic on-target response of the CRISPRa module could be mitigated by increasing dCas9 levels. To this end, we used the same construct for a single CRISPRa module as used in Fig. 1, but with variable dCas9 promoter strength (Fig. 5c). We observed minimal change to the biphasic response of RFP to scRNA_RFP_ level as the dCas9 expression was increased (Fig. 5d). It follows that increasing dCas9 level does not alter the biphasic on-target response of a CRISPRa module.

Next, we investigated whether increasing dCas9 level could at least mitigate the coupling between two independent CRISPRa modules. To this end, we assembled genetic constructs comprising two CRISPRa modules in which we could vary the promoter strength of dCas9 (Fig. 5e). Although increased dCas9 level reduced the drop of RFP from 85% to 45% when the scRNA_GFP_ level was increased, it could not decouple the two modules (Fig. 5f and SI Fig. 18). Further increasing dCas9 expression beyond the level obtained with the strong pCas promoter showed growth defects (SI Fig. 19), indicating that the strongest promoter used in our experiments (Fig. 5) achieves the maximal level of dCas9 that is tolerated by the cell. Therefore, increasing dCas9 level can reduce but not remove the coupling between the two modules.

We also investigated whether we could increase the expression of RBP-AD beyond the level expressed in our construct (Fig. 5). However, substantial cell death was observed beyond this point (SI Fig. 19), indicating that the promoter used for RBP-AD in our experiments produces the maximal level of RBP-AD that is tolerated by the cell.

Taken together, our results show that it is not possible to remove the observed coupling between the two modules by increasing the resource levels since coupling remains at the maximal levels tolerated by the cell.

### Optimizing the response of a CRISPRa module by expression of a competitor scRNA

Since the output of a CRISPRa module decreases following the expression of a competitor scRNA (Fig. 3d), we investigated whether we could take advantage of this fact to decrease the leakiness of a CRISPRa module and to thus increase its dynamic range. To this end, we assembled a genetic system containing two CRISPRa modules such that the scRNA of Module 2 can be regarded as a competitor scRNA for Module 1. We interrogated the model to determine how changing different aspects of the competitor scRNA, including its DNA binding target (see SI Note 3.2), could affect the input/output response of Module 1 (Fig. 6a). According to the model, adding a competitor scRNA lacking a DNA target sequence is the most effective approach to decrease the basal output expression of a CRISPRa module while minimizing the output reduction for large inputs (Fig. 6b). In fact, adding a competitor scRNA (with or without a DNA target) reduces the Module 1 RFP output level significantly due to competition for resources (Fig. 3e). On the other hand, the RFP output of Module 1 for high input scRNA (large DAPG) is higher when the competitor scRNA does not bind to a target site because the binding with DNA stabilizes the CRISPR complex and therefore sequesters more resources when compared to an scRNA that does not bind to DNA.

**Figure 6.**
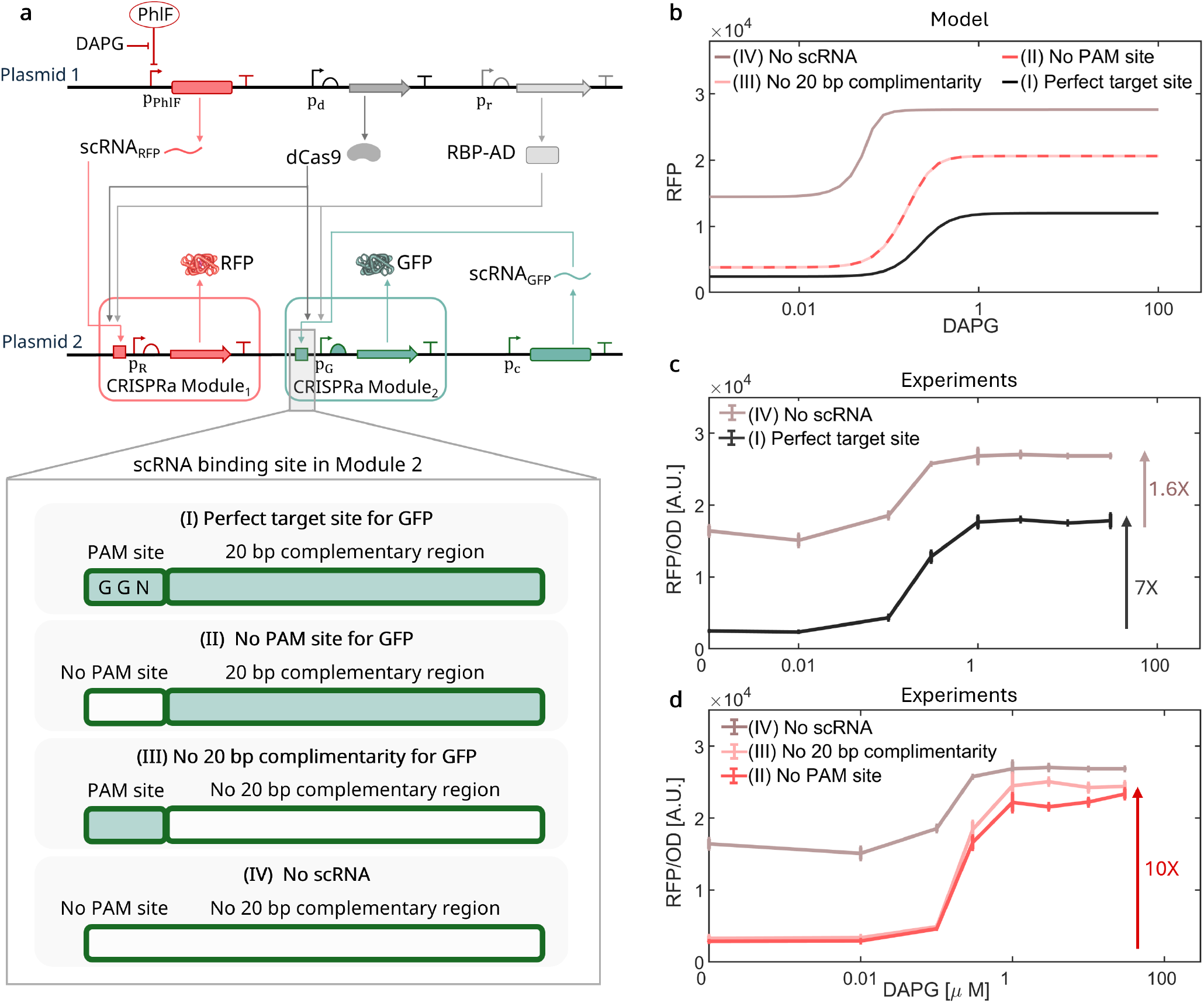
Optimizing the input-output response of a CRISPRa module by expressing a competitor scRNA. (a) Genetic construct for two CRISPRa modules with scRNA_RFP_ activating RFP and scRNA_GFP_ activating GFP. The DNA binding site for scRNA_GFP_ is changed as indicated: (I) has both perfect binding site to region in the scRNA_GFP_ and a PAM site, (II) has no PAM site, (III) has no 20 bp complementarity region, and (IV) has no competitor scRNA_GFP_. Sequences for the target site variations are in SI Note 4.1. (b) Numerical simulation results showing RFP with increasing DAPG for the four cases in (a) using the ODE model in SI Note 3.2. Here, (II) and (III) are modeled as not having a target gene since the model does not have separate PAM and 20bp complementarity sites. (c) Response of RFP to DAPG in the presence of the scRNA_GFP_ binding site (I) and (IV). (d) Response of RFP to DAPG in the presence of the scRNA_GFP_ binding site (II) and (III). Temporal data is provided in SI Fig. **20**.

Our experiments confirm the model predictions (Figs. 6c,d). Specifically, in the absence of a competitor scRNA, the Module 1 RFP output level shows a 1.6-fold activation as the scRNA_RFP_ is increased via DAPG. The addition of a competitor scRNA with a DNA target site increases the fold activation to 7-fold, but also reduces the RFP levels at maximum DAPG induction (Fig. 6c). In order to increase the fold activation while maintaining the maximum activation level, we introduced a competitor scRNA without a target site (either lacking the PAM site or the complementary DNA target sequence). This leads to a fold activation of approximately 10 with diminished decrease in the maximum activation level (Fig. 6d). Therefore, the addition of a competitor scRNA without a DNA target site can be used as an effective strategy to optimize the on-target response of a CRISPRa module. Changing the level of dCas9 does not change these trends (see SI Fig. **21**).

## Discussion

In this paper, we investigated the effects of competition for dCas9 and RBP-AD in CRISPRa systems. We discovered that the on-target response of a CRISPRa module is biphasic instead of being monotonically increasing (Fig. 1) and introduced the double-diamond reaction topology found in CRISPRa as a plausible mechanism for it (Fig. 2). We further showed that two CRISPRa modules become coupled due to competition for dCas9 and RBP-AD proteins (Fig. 3) and that both of these two resources are responsible for the observed competition effects (Fig. 4). In turn, increasing the resource level to the maximal value tolerated by the cell could not eliminate competition effects (Fig. 5). Although regarded as a negative for the engineering of modular systems, we leveraged competition to optimize the input/output response of a CRISPRa module without altering the genetic sequence of the module components (Fig. 6).

As the effects of competition could not be removed by increasing the dCas9 or RBP-AD levels, restoring modularity and the desired increasing on-target CRISPRa responses will likely require the design of feedback controllers that maintain both dCas9 and RBP-AD resources at constant levels despite changing demands by scRNAs. These controllers have been designed already for dCas9 and have demonstrated that competition in CRISPRi systems can be neutralized this way ^22^. For CRISPRa, however, since competition effects arise also due to RBP-AD sharing (Fig. 4), the introduction of an RBP-AD regulator will likely be required to quench the effects of competition.

More broadly, our results indicate that competition for dCas9 and RBP-AD plays a key role in the emergent function of CRISPRa networks by coupling theoretically orthogonal modules and by making a supposedly monotonic CRISPRa response biphasic. Our mathematical model establishes a predictive tool that captures these emergent effects and can be used for the design of CRISPRa networks that behave as intended despite the lack of modularity. These models could further be used to determine how competition affects the outcomes of studies that use multiple scRNAs and gRNAs concurrently, such as in genome-wide screening, where orthogonality is assumed but fails in practice.

## Methods

### Bacterial strain and media

The bacterial strain *E. coli* NEB10B (NEB, C3019I) grown in LB Broth Lennox (240230) with appropriate antibiotics were used for genetic circuit construction. The sequence-verified plasmids are co-transformed or retransformed to *E. coli* Marionette strain ^29^ for all presented results. The experiment’s growth medium is an M9 minimal medium composed of M9 salts (1X), 0.4% glucose (SIGMA-ALDRICH, 49159), 0.2% casamino acid (VWR, TS61204-5000), and 1mM thiamine hydrochloride (SIGMA-ALDRICH, T4625-25G) supplemented by appropriate antibiotics. The final concentrations of antibiotics, ampicillin (SIGMA-ALDRICH, A0166-5G), kanamycin (SIGMA-ALDRICH, K1377-5G), and chloramphenicol (SIGMA-ALDRICH, C0378-5G) are 100, 50, and 34 *µ*g mL^−1^, respectively. 2,4-Diacetylphloroglucinol or DAPG (Cayman Chemical, 16345) is used as the inducer.

### Genetic circuit construction

The genetic circuit construction was based on Gibson assembly methods ^31^ using pCK005.6 and pJF143.J3 from Addgene ^11^ as the backbone. DNA oligos and genes were ordered using the Oligo Rapid Service from Azenta Life Sciences. DNA fragments to be assembled were amplified by PCR using Phusion High-Fidelity PCR Master Mix with GC Buffer (NEB, M0532S). The PCR mix underwent gel electrophoresis for purification for 30 minutes at 110V. Zymoclean Gel DNA Recovery Kit (Zymo Research, D4002) was used for gel extraction. Later, the plasmid is assembled with Gibson assembly protocol using NEBuilder HiFi DNA Assembly Master Mix (NEB, E2621S). Assembled DNA was transformed into NEB10B competent cells prepared by the CCMB80 buffer (TekNova, C3132). The transformed cells are then plated on Agar plates with 1.2% of Difco Agar and 2% of LB broth Lennox and appropriate antibiotics and grown overnight in an incubator at 37^*o*^C. Overnight cultures of singular colonies picked from the agar plates are grown in culture tubes with LB broth Lennox at 37^*o*^C. Plasmid DNA is then extracted from the overnight cultures using the plasmid miniprep-classic kit (Zymo Research, D4015). DNA sequencing is outsourced and is performed by Plasmidsaurus: Standard Low Concentration Plasmid sequencing. The lists of plasmids and genetic parts are listed in SI Notes 1 and 4, respectively.

### Plate reader experiments

The glycerol stocks (stored at -80^*o*^C) are used to prepare overnight cultures. These cultures are grown in M9 media supplemented with appropriate antibiotics at 37 ^*o*^C, shaking at 220 rpm in a horizontal orbiting shaker for 6-8 hours in 15 mL culture tubes (VWR, 60818-667). These overnight cultures are diluted to an optical density at 600nm (OD600) measurement of 0.01 (after subtracting background signal from media) in 200 uL growth medium per well in a transparent 96-well plate with a flat bottom (FalconR 96-Well Cell Culture Plates, Corning, 15705-066) with appropriate antibiotics and inducers. The 96-well plate was incubated at 30^*o*^C in a Tecan Infinite 200 PRO microplate reader in static condition and was shaken at a fast speed for 5 seconds right before OD and fluorescence measurements. The sampling interval was 5 min. Excitation and emission wavelengths to monitor RFP fluorescence were 584 and 619 nm, for GFP fluorescence were 485 and 530 nm, and for YFP fluorescence were 488 and 530 nm, respectively. To ensure continued exponential growth, cell cultures were diluted with fresh growth medium (with antibiotics and inducers) to OD600 of 0.01 when OD600 approaches 0.08 at the end of one batch. Multiple batches were conducted with the total experiment time of up to 36 hours until gene expression reached a steady state. The reported steady-state fluorescent values are normalized with the OD600 and are reported when each culture is at an OD600 of 0.06 in the final batch.

### Statistics and reproducibility

Statistical significance was calculated using two-tailed two sample t-tests with the MATLAB 2022 b Statistics and Machine Learning Toolbox. To ensure reproducibility, experiments were performed using 3 biological replicates.

## Acknowledgements

This work was supported by NSF CCF-FET Award 2007674.

## References

1. P. Mali, J. Aach, P. B. Stranges, K. M. Esvelt, M. Moosburner, S. Kosuri, L. Yang, and G. M. Church, “Cas9 transcriptional activators for target specificity screening and paired nickases for cooperative genome engineering,” Nature Biotechnology, vol. 31, no. 9, pp. 833–838, 2013.

2. F. J. Mojica, G. Juez, and F. Rodriguez-Valera, “Transcription at different salinities of haloferax mediterranei sequences adjacent to partially modified Psti sites,” Molecular Microbiol., vol. 9, no. 3, pp. 613–621, 1993.

3. E. S. Lander, “The heroes of CRISPR,” Cell, vol. 164, no. 1, pp. 18–28, 2016.

4. A. Bolotin, B. Quinquis, A. Sorokin, and S. D. Ehrlich, “Clustered regularly interspaced short palindrome repeats (CRISPRs) have spacers of extrachromosomal origin,” Microbiology, vol. 151, no. 8, pp. 2551–2561, 2005.

5. E. Deltcheva, K. Chylinski, C. M. Sharma, K. Gonzales, Y. Chao, Z. A. Pirzada, M. R. Eckert, J. Vogel, and E. Charpentier, “CRISPR RNA maturation by trans-encoded small RNA and host factor RNase iii,” Nature, vol. 471, no. 7340, pp. 602–607, 2011.

6. G. Gasiunas, R. Barrangou, P. Horvath, and V. Siksnys, “Cas9–crRNA ribonucleoprotein complex mediates specific DNA cleavage for adaptive immunity in bacteria,” Proceedings of the National Academy of Sciences, vol. 109, no. 39, pp. E2579–E2586, 2012.

7. M. Jinek, K. Chylinski, I. Fonfara, M. Hauer, J. A. Doudna, and E. Charpentier, “A programmable dual-RNA–guided DNA endonuclease in adaptive bacterial immunity,” Science, vol. 337, no. 6096, pp. 816–821, 2012.

8. L. Cong, F. A. Ran, D. Cox, S. Lin, R. Barretto, N. Habib, P. D. Hsu, X. Wu, W. Jiang, L. A. Marraffini, et al., “Multiplex genome engineering using CRISPR/Cas systems,” Science, vol. 339, no. 6121, pp. 819–823, 2013.

9. P. Mali, L. Yang, K. M. Esvelt, J. Aach, M. Guell, J. E. DiCarlo, J. E. Norville, and G. M. Church, “RNA-guided human genome engineering via Cas9,” Science, vol. 339, no. 6121, pp. 823–826, 2013.

10. A. A. Dominguez, W. A. Lim, and L. S. Qi, “Beyond editing: repurposing CRISPR–Cas9 for precision genome regulation and interrogation,” Nature Reviews Molecular Cell Biology, vol. 17, no. 1, pp. 5–15, 2016.

11. J. Fontana, C. Dong, C. Kiattisewee, V. P. Chavali, B. I. Tickman, J. M. Carothers, and J. G. Zalatan, “Effective CRISPRa-mediated control of gene expression in bacteria must overcome strict target site requirements,” Nature Communications, vol. 11, no. 1, p. 1618, 2020.

12. C. Dong, J. Fontana, A. Patel, J. M. Carothers, and J. G. Zalatan, “Synthetic CRISPR-Cas gene activators for transcriptional reprogramming in bacteria,” Nature Communications, vol. 9, no. 1, pp. 1–11, 2018.

13. C. Bock, P. Datlinger, F. Chardon, M. A. Coelho, M. B. Dong, K. A. Lawson, T. Lu, L. Maroc, T. M. Norman, B. Song, et al., “High-content CRISPR screening,” Nature Reviews Methods Primers, vol. 2, no. 1, pp. 1–23, 2022.

14. S. Bodapati, T. P. Daley, X. Lin, J. Zou, and L. S. Qi, “A benchmark of algorithms for the analysis of pooled CRISPR screens,” Genome Biology, vol. 21, pp. 1–13, 2020.

15. T. Clark, M. A. Waller, L. Loo, C. L. Moreno, C. E. Denes, and G. G. Neely, “CRISPR activation screens: navigating technologies and applications,” Trends in Biotechnology, 2024.

16. J. Fontana, D. Sparkman-Yager, J. G. Zalatan, and J. M. Carothers, “Challenges and opportunities with CRISPR activation in bacteria for data-driven metabolic engineering,” Current Opinion in Biotechnology, vol. 64, pp. 190–198, 2020.

17. J. Fontana, D. Sparkman-Yager, I. Faulkner, R. Cardiff, C. Kiattisewee, A. Walls, T. G. Primo, P. C. Kinnunen, H. Garcia Martin, J. G. Zalatan, et al., “Guide RNA structure design enables combinatorial CRISPRa programs for biosynthetic profiling,” bioRxiv, pp. 2023–11, 2023.

18. W. M. Shaw, L. Studená, K. Roy, P. Hapeta, N. S. McCarty, A. E. Graham, T. Ellis, and R. Ledesma-Amaro, “Inducible expression of large gRNA arrays for multiplexed CRISPRai applications,” Nature Communications, vol. 13, no. 1, p. 4984, 2022.

19. M. Kampmann, “CRISPRi and CRISPRa screens in mammalian cells for precision biology and medicine,” ACS Chemical Biology, vol. 13, no. 2, pp. 406–416, 2018.

20. A. C. Bester, J. D. Lee, A. Chavez, Y.-R. Lee, D. Nachmani, S. Vora, J. Victor, M. Sauvageau, E. Monteleone, J. L. Rinn, et al., “An integrated genome-wide CRISPRa approach to functionalize lncRNAs in drug resistance,” Cell, vol. 173, no. 3, pp. 649–664, 2018.

21. P.-Y. Chen, Y. Qian, and D. Del Vecchio, “A model for resource competition in CRISPR-mediated gene repression,” in 2018 IEEE Conference on Decision and Control (CDC), pp. 4333–4338, IEEE, 2018.

22. H.-H. Huang, M. Bellato, Y. Qian, P. Cárdenas, L. Pasotti, P. Magni, and D. Del Vecchio, “dCas9 regulator to neutralize competition in CRISPRi circuits,” Nature Communications, vol. 12, no. 1, p. 1692, 2021.

23. D. A. Anderson and C. A. Voigt, “Competitive dcas9 binding as a mechanism for transcriptional control,” Molecular Systems Biology, vol. 17, no. 11, p. e10512, 2021.

24. S. Zhang and C. A. Voigt, “Engineered dCas9 with reduced toxicity in bacteria: implications for genetic circuit design,” Nucleic Acids Research, vol. 46, no. 20, pp. 11115–11125, 2018.

25. K. Manoj and D. Del Vecchio, “Emergent interactions due to resource competition in CRISPR-mediated genetic activation circuits,” in 2022 IEEE 61st Conference on Decision and Control (CDC), pp. 1300–1305, IEEE, 2022.

26. S. Clamons and R. Murray, “Modeling predicts that CRISPR-based activators, unlike CRISPR-based repressors, scale well with increasing gRNA competition and dCas9 bottlenecking,” bioRxiv, p. 719278, 2019.

27. B. I. Tickman, D. A. Burbano, V. P. Chavali, C. Kiattisewee, J. Fontana, A. Khakimzhan, V. Noireaux, J. G. Zalatan, and J. M. Carothers, “Multi-layer CRISPRa/i circuits for dynamic genetic programs in cell-free and bacterial systems,” Cell Systems, vol. 13, no. 3, pp. 215–229, 2022.

28. S. Kosuri, D. B. Goodman, G. Cambray, V. K. Mutalik, Y. Gao, A. P. Arkin, D. Endy, and G. M. Church, “Composability of regulatory sequences controlling transcription and translation in Escherichia coli,” Proceedings of the National Academy of Sciences, vol. 110, no. 34, pp. 14024–14029, 2013.

29. A. J. Meyer, T. H. Segall-Shapiro, E. Glassey, J. Zhang, and C. A. Voigt, “Escherichia coli “Marionette” strains with 12 highly optimized small-molecule sensors,” Nature Chemical Biology, vol. 15, no. 2, pp. 196–204, 2019.

30. A. Gyorgy, J. I. Jiménez, J. Yazbek, H.-H. Huang, H. Chung, R. Weiss, and D. Del Vecchio, “Isocost lines describe the cellular economy of genetic circuits,” Biophysical journal, vol. 109, no. 3, pp. 639–646, 2015.

31. D. G. Gibson, L. Young, R.-Y. Chuang, J. C. Venter, C. A. Hutchison III, and H. O. Smith, “Enzymatic assembly of dna molecules up to several hundred kilobases,” Nature methods, vol. 6, no. 5, pp. 343–345, 2009.

