## Supplementary Information for "Resource competition shapes CRISPR-mediated gene activation"

#### 1 Supplementary Note 1: Plasmid maps and constituents

In this section, we provide details on the different plasmid maps used in the study along with details for the genetic constructs. We mainly co-transform two plasmids into the *E. coli* Marionette strain[1] for all presented results. Plasmid 1 typically contains dCas9, RBP-AD and an scRNA while plasmid 2 contains all reporter proteins and the competitor scRNA. The specific sequences for all the components are in SI Section 4.

| Figure Name | Plasmids used | Component Varied |
|---|---|---|
| Main Fig. 1 d | Plasmid 1 and Plasmid 2a | scRNA promoters $p_{s1}$ and $p_{s2}$ |
| Main Fig. 3 c | Plasmid 1 and Plasmid 2b | DAPG induced scRNA$_{RFP}$ |
| Main Fig. 3 d | Plasmid 1 and Plasmid 2b | scRNA$_{GFP}$ promoter $p_C$ |
| Main Fig. 3 e | Plasmid 1 and Plasmid 2b | scRNA$_{GFP}$ as shown in SI section 4.2 |
| Main Fig. 4 b | Plasmid 1 and Plasmid 2b | scRNA$_{GFP}$ as shown in SI section 4.2 |
| Main Fig. 5 b | Plasmid 1 and Plasmid 2c | promoters for dCas9 $p_d$ and DAPG induced scRNA$_{RFP}$ |
| Main Fig. 5 d | Plasmid 1 and Plasmid 2a | promoters for dCas9 $p_d$ and scRNA $p_{s1}$ and $p_{s2}$ |
| Main Fig. 5 f | Plasmid 1 and Plasmid 2b | promoters for dCas9 $p_d$ with and without scRNA$_{GFP}$ as shown in SI section 4.2 |
| Main Fig. 6 c | Plasmid 1 and Plasmid 2b | DAPG induced scRNA$_{RFP}$ with and without scRNA$_{GFP}$ (see SI section 4.1, 4.2) |
| Main Fig. 6 d | Plasmid 1 and Plasmid 2b | DAPG induced scRNA$_{RFP}$ and varying GFP binding sites (see SI section 4.1) |

**Table 1.** List of plasmids used in the Main Manuscript along with the components that are varied in each figure.

##### 1.1 Plasmid 1

Plasmid 1 consists of dCas9, RBP-AD, and scRNA_RFP (scRNA designed to target the RFP gene) with varying promoters for the scRNA and dCas9 as given in Table 2 (SI Fig. 1a). Plasmid 1 is on a p15A vector (20 copies per cell[2]) and is Kanamycin antibiotic resistant.

##### 1.2 Plasmid 2

Three variations of plasmid 2 have been used in the entire study. All three variants have Ampicillin antibiotic resistance.

###### *Plasmid 2a*

Plasmid 2a has been used to study the biphasic response of a single CRISPRa module (Main Manuscript Fig. 1d and 5d) and is on a PSC101(E93G) vector, 84 copies per cell[2](SI Fig. 1b). It consists of the target RFP gene constitutively expressed through the minimal promoter J117 and RBS_RFP. The J306 binding site for the scRNA$_{RFP}$ is placed -81 bp upstream of the TSS of the RFP gene. The additional scRNA (that binds to the J306 binding site) is constitutively expressed with varying promoter strengths shown in Table 2.

###### *Plasmid 2b*

Plasmid 2b has been used to study the behavior of two CRISPRa modules (Main Manuscript Fig. 3, 4, 5f and 6) and is on a PSC101 vector, 5 copies per cell[2](SI Fig. 1c). It consists of the target RFP gene (the output of CRISPRa Module 1) constitutively expressed through the minimal promoter J117 and RBS_RFP. The J306 binding site for the scRNA$_{RFP}$ is placed -81 bp upstream of the TSS of the RFP gene. It also hosts the input scRNA (that binds to the J108 binding site) and the output GFP (with the J108 binding site) of CRISPRa module 2. The target GFP site also uses the minimal promoter J117 with RBS_GFP.

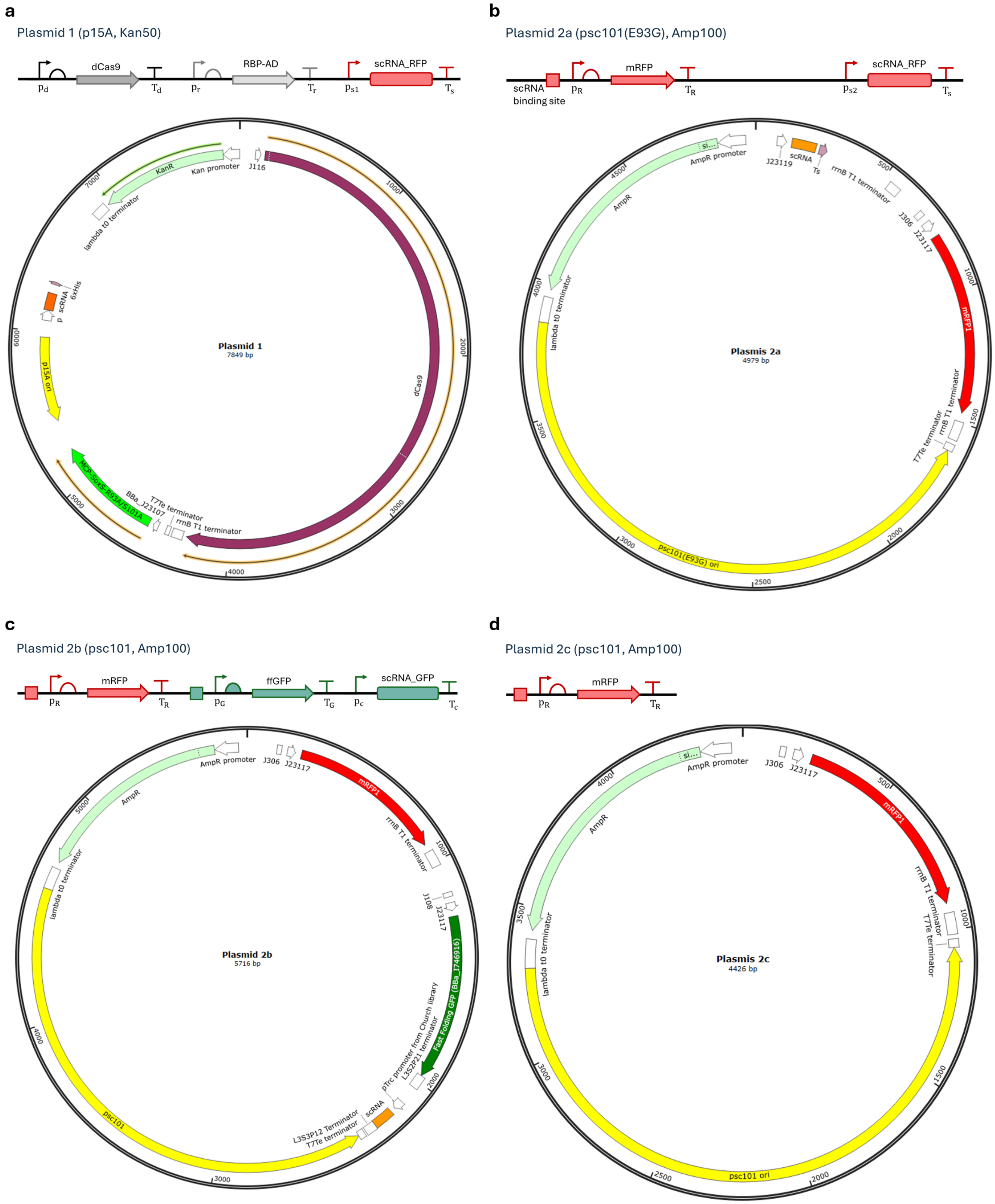


**SI Figure 1.** Genetic maps used in constructs listed in Table 1. The plasmids shown are (a) Plasmid 1, (b) Plasmid 2a, (c) Plasmid 2b, and (d) Plasmid 2c.

**Plasmid 2c**
Plasmid 2c has been used to study the behavior of a single CRISPRa module (Main Manuscript Fig. 5b) and is on a PSC101 vector, 5 copies per cell[2](SI Fig. 1d). It hosts the target RFP gene (the output of CRISPRa Module 1), constitutively expressed through the minimal promoter J117 and RBS_RFP. The J306 binding site for the $scRNA_{RFP}$ is placed -81 bp upstream of the TSS of the RFP gene.

##### 1.3 Promoters, RBS and Terminators in each cassette

| Figure Name | dCas9 promoter $p_d$ | $scRNA_{RFP}$ promoter $p_{s1}$ | $scRNA_{RFP}$ promoter $p_{s2}$ | $scRNA_{GFP}$ promoter $p_C$ |
|---|---|---|---|---|
| Main Fig. 1 d | BBa_J23116 | P1 = pTac (no IPTG)<br>P2 = pTac (with IPTG)<br>P3 = pPhlF | P4 = pTac (no IPTG)<br>P5 = BBa_J23110<br>P6 = BBa_J23119 | NA |
| Main Fig. 3 c | BBa_J23116 | pPhlF | NA | pTrc |
| Main Fig. 3 d | BBa_J23116 | pPhlF | NA | Low - BBa_J23114<br>Medium - LacUV5<br>High - pTrc |
| Main Fig. 3 e | BBa_J23116 | pPhlF | NA | pTrc |
| Main Fig. 4 b | BBa_J23116 | pPhlF | NA | pTrc |
| Main Fig. 5 b | BBa_J23103<br>BBa_J23117<br>BBa_J23115<br>BBa_J23116<br>BBa_J23105<br>Sp.pCas9* | pPhlF | NA | NA |
| Main Fig. 5 d | BBa_J23116<br>Sp.pCas9* | P1 = pTac (no IPTG)<br>P2 = pTac (with IPTG)<br>P3 = pPhlF | P4 = pTac (no IPTG)<br>P5 = BBa_J23110<br>P6 = BBa_J23119 | NA |
| Main Fig. 5 f | BBa_J23116<br>Sp.pCas9* | $p_s$ = pPhlF | NA | pTrc |
| Main Fig. 6 c | BBa_J23116 | pPhlF | NA | pTrc |
| Main Fig. 6 d | BBa_J23116 | pPhlF | NA | pTrc |

**Table 2.** List of promoters varied in each figure in the Main Manuscript. Note: *Sp. pCas9* promoter is the endogenous promoter from the *S. pyogenes* dCas9 protein and is always combined with its endogenous RBS.

| Protein/ RNA species | Promoter | RBS | Terminator |
|---|---|---|---|
| dCas9 | $p_d$ varied (Table 2) | BBa_B0034 | BBa_B0015 |
| RBP-AD | $p_r$ = BBa_J23107 | BBa_J34801 | BBa_B1002 |
| $scRNA_{RFP}$ | $p_{s1}$ and $p_{s2}$ varied (Table 2) | NA | BBa_K1893035 |
| mRFP1 | $p_R$ = BBa_J23117 | RBS_RFP | rrnB T1 |
| $scRNA_{GFP}$ | $p_c$ varied (Table 2) | NA | L3S3P12 + T7Te |
| FF GFP (BBa_I746916) | $p_G$ = BBa_J23117 | RBS_GFP | L3S2P21 |

**Table 3.** List of cassettes with their promoters. RBS and terminators for the all figures in the Main Manuscript

### 2 Supplementary Note 2: Extended Data and Figures

#### 2.1 Promoter strength characterization for scRNA in Main Manuscript Figure 1

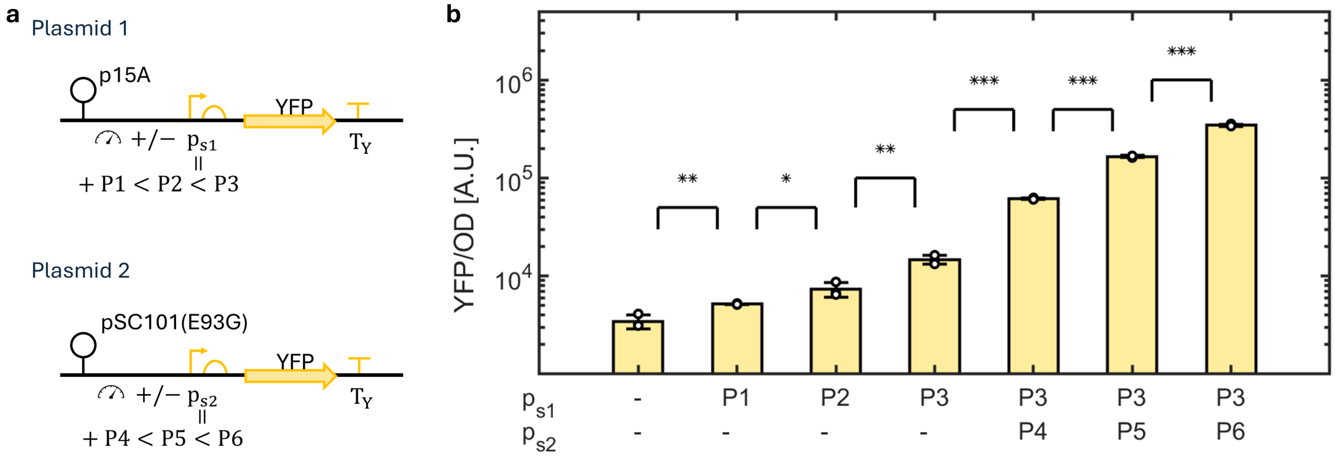


**SI Figure 2.** Promoter strength characterization for the various promoters used to vary the amount of scRNA to observe the biphasic on-target response in Main Manuscript Fig. 1. (a) Genetic construct producing YFP with varying promoters, used to quantify the promoter strengths of scRNAs in Main Manuscript Fig. 1(d). Towards this, YFP is placed on two plasmids with the same promoters ($p_{s1}$ and $p_{s2}$), origin of replications, and antibiotic resistances as the scRNAs in Main Manuscript Fig. 1(b). (b) Bar chart representation of YFP showing the relative increase in the production while using the combinations of promoters implemented in (a).

#### 2.2 Temporal data for each data point in Main Manuscript Figure 1(d)

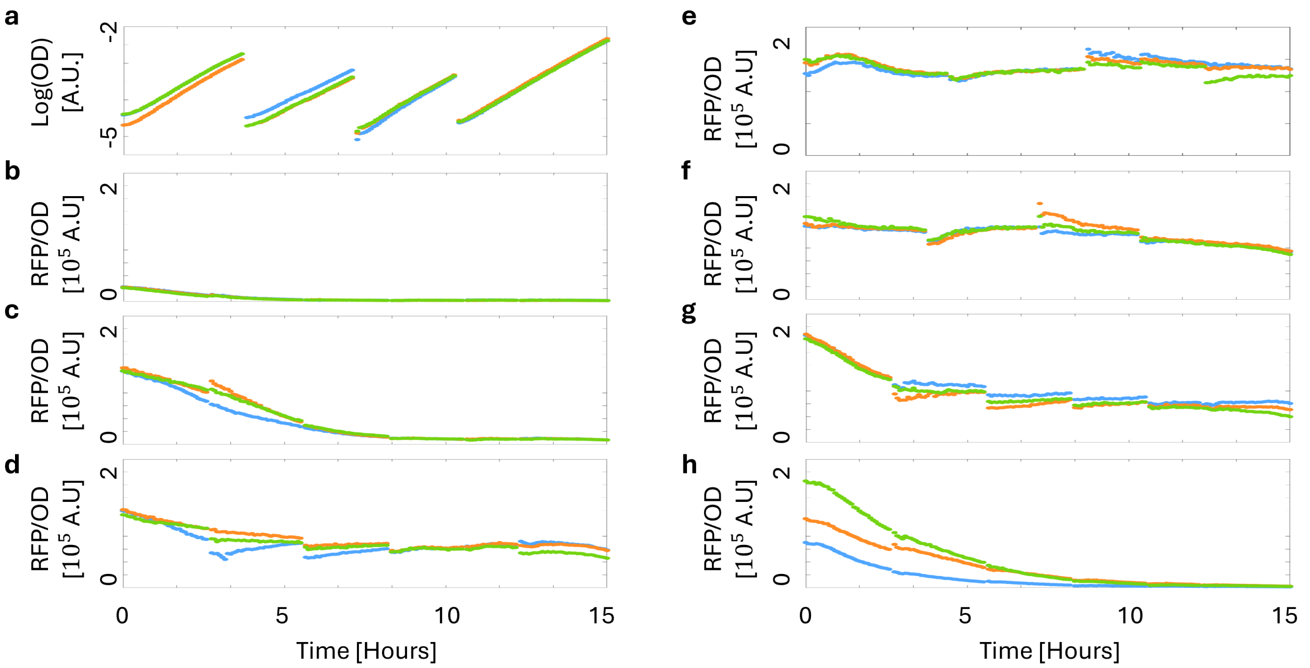


**SI Figure 3.** Time Series data for each data point in Main Manuscript Fig. 1(d) using the genetic construct shown in Main Manuscript Fig. 1(b). (a) Temporal data for log(measured OD - background OD) for four batches spanning a total of 15 hours for the scRNA promoter combinations of $p_{s1} = -$ and $p_{s2} = -$. The cells are maintained in the exponential phase by serial dilution as explained in the methods. (b - h) Temporal evolution of RFP for different levels of input scRNA. Here the corresponding scRNA promoter combinations are (b) $p_{s1} = -$ and $p_{s2} = -$, (c) $p_{s1} =$ P1 and $p_{s2} = -$, (d) $p_{s1} =$ P2 and $p_{s2} = -$, (e) $p_{s1} =$ P3 and $p_{s2} = -$, (f) $p_{s1} =$ P3 and $p_{s2} =$ P4, (g) $p_{s1} =$ P3 and $p_{s2} =$ P5, and (h) $p_{s1} =$ P3 and $p_{s2} =$ P6, respectively. The steady states used in the Main Manuscript correspond to the time point when the measured OD reaches 0.14 (approximately - 2.8 in a) in the final fourth batch.

##### 2.3 Variation of each molecule in the double diamond reaction diagram as the input scRNA is increased

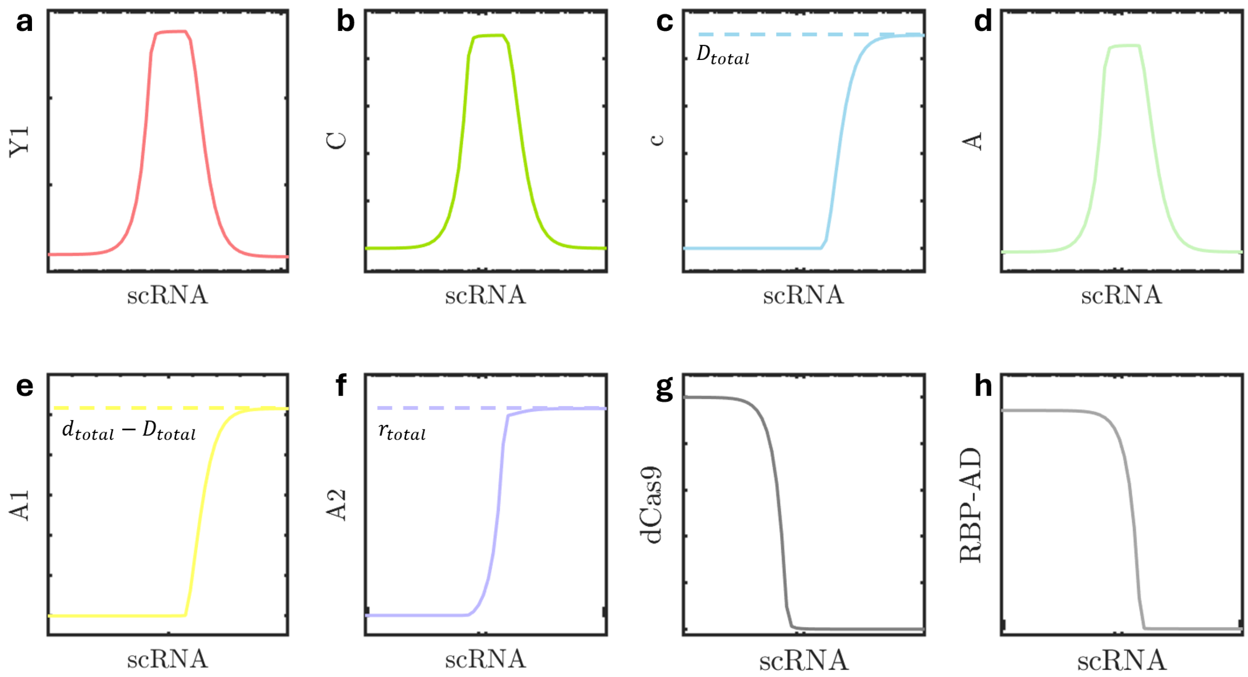


**SI Figure 4.** Line plots of levels of each complex and resources in a CRISPRa module as the input scRNA is increased. The line plots correspond to (a) output protein ($Y$), (b) transcriptionally active complex ($C$), (c) intermediate scRNA-dCas9-DNA complex ($c$), (d) activator complex ($A$), (e) intermediate scRNA-dCas9 complex ($A_1$), (f) intermediate scRNA-RBP-AD complex ($A_2$), (g) free dCas9 ($d$), and (h) free RBP-AD ($r$). From these plots, we see that as scRNA is increased, the dCas9 resources are sequestered to form complexes $c$ and $A_1$ while RBP-AD resources are sequestered to form the complex $A_2$. This causes a decrease in the amounts of $C$, which in turn leads to a biphasic response in $Y$.

#### 2.4 Promoter characterizations for low, medium, and high promoters in Main Manuscript Figure 3(d)

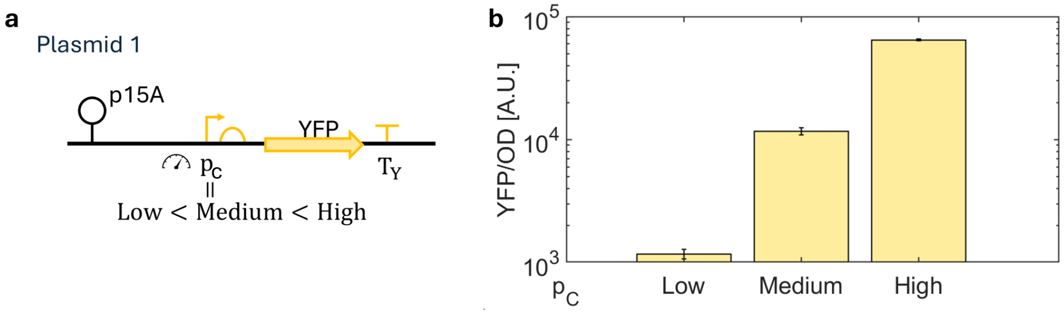


**SI Figure 5.** Promoter strength characterization for the promoters for the scRNA$_{GFP}$ used in Main Manuscript Fig. 3(d). (a) Genetic construct producing YFP with low, medium and high promoters used to quantify the promoter strengths of scRNA$_{GFP}$ in Main Manuscript Fig. 3(d). The specific promoters are low (BBa_J23114), medium (LacUV5), and high (pTrc), see Table 5 for sequences. (b) Bar chart representation of YFP showing the relative strengths of the promoters used.

#### 2.5 Temporal data for each data point in Main Manuscript Figure 3

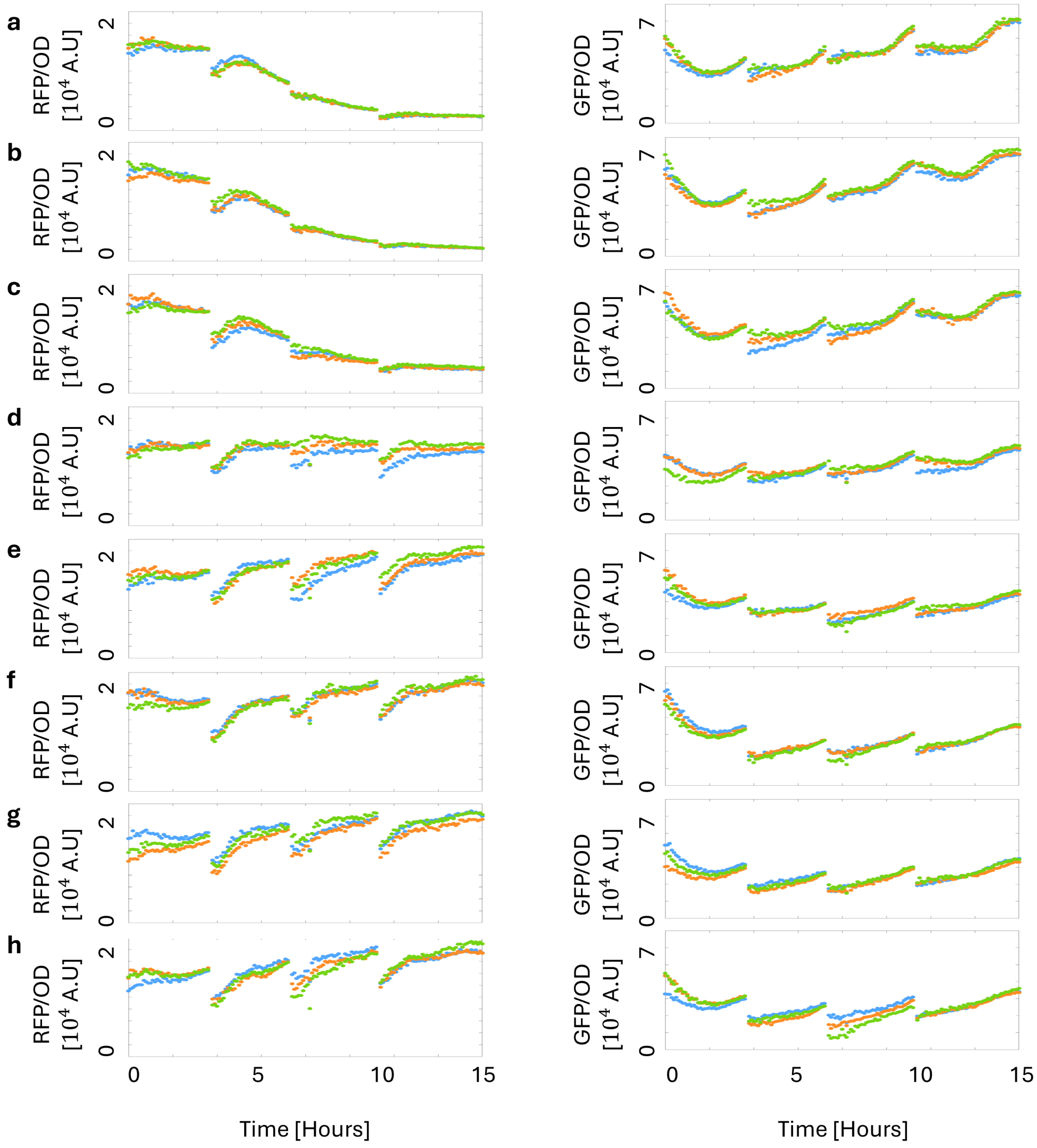


**SI Figure 6.** Time Series data for each data point in Main Manuscript Fig. 3(c) using the genetic construct shown in Main Manuscript Fig. 3(b). (a-h) Temporal evolution of (Left) RFP and (Right) GFP for different levels of DAPG. Here the DAPG levels are (a) $0\mu M$, (b) $0.01\mu M$, (c) $0.1\mu M$, (d) $0.3\mu M$, (e) $1\mu M$, (f) $3\mu M$, (g) $10\mu M$, and (h) $30\mu M$, respectively. The steady states used in the Main Manuscript correspond to the time point when the measured OD reaches 0.14 in the final fourth batch.

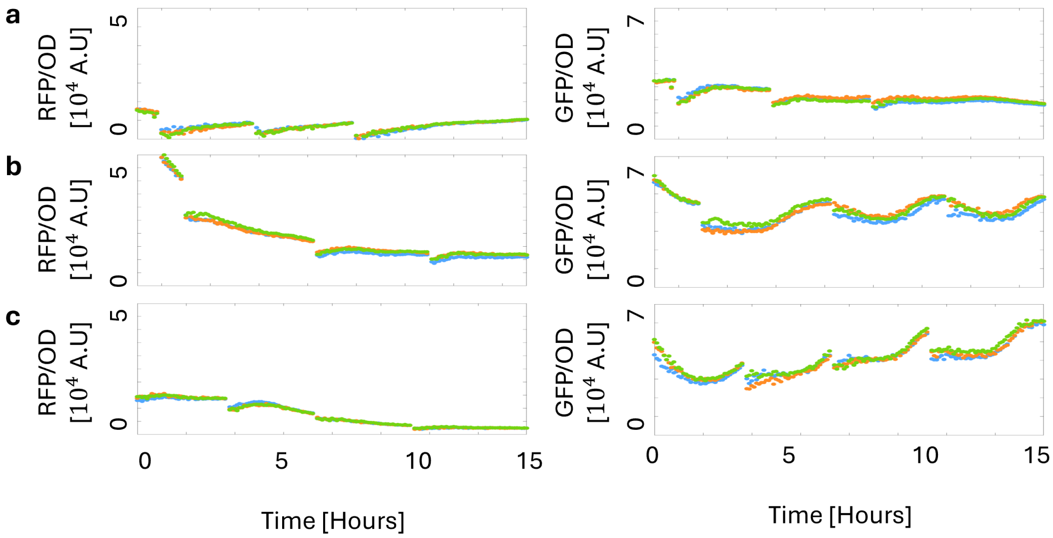


**SI Figure 7.** Time Series data for each data point in Main Manuscript Fig. 3(d) using the genetic construct shown in Main Manuscript Fig. 3(b). (a-c) Temporal evolution of (Left) RFP and (Right) GFP for different promoters for scRNA$_{GFP}$. Here the promoter strengths are (a) low, (b) medium, and (c) high, respectively. The steady states used in the Main Manuscript correspond to the time point when the measured OD reaches 0.14 in the final fourth batch.

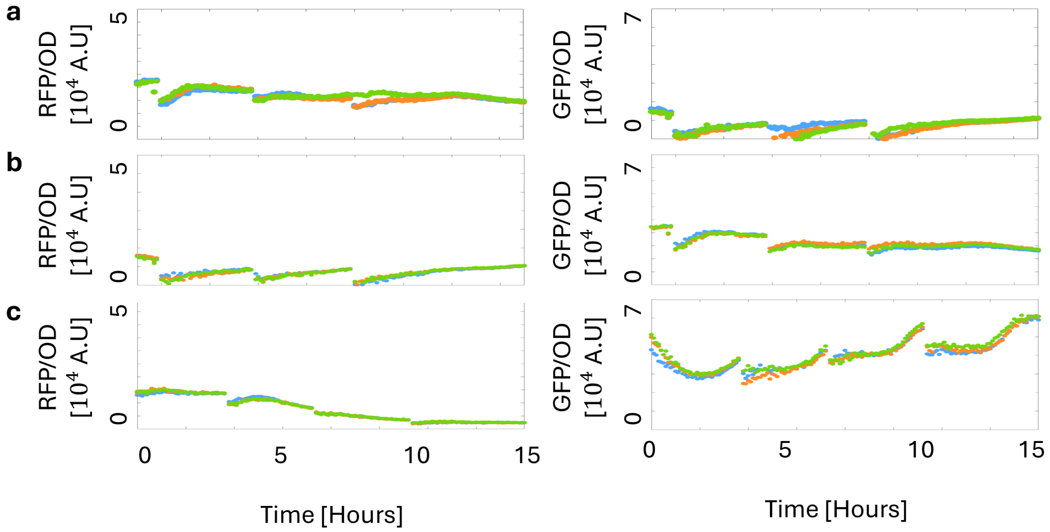


**SI Figure 8.** Time Series data for each data point in Main Manuscript Fig. 3(e) using the genetic construct shown in Main Manuscript Fig. 3(b). (a-c) Temporal evolution of (Left) RFP and (Right) GFP for different scRNA$_{GFP}$. Here the different scRNA$_{GFP}$ configurations are (a) no scRNA$_{GFP}$, (b) scrambled scRNA$_{GFP}$, and (c) with perfect scRNA$_{GFP}$, respectively. The steady states used in the Main Manuscript correspond to the time point when the measured OD reaches 0.14 in the final fourth batch.

##### 2.6 Coupling between GFP and RFP for low and medium scRNA_GFP

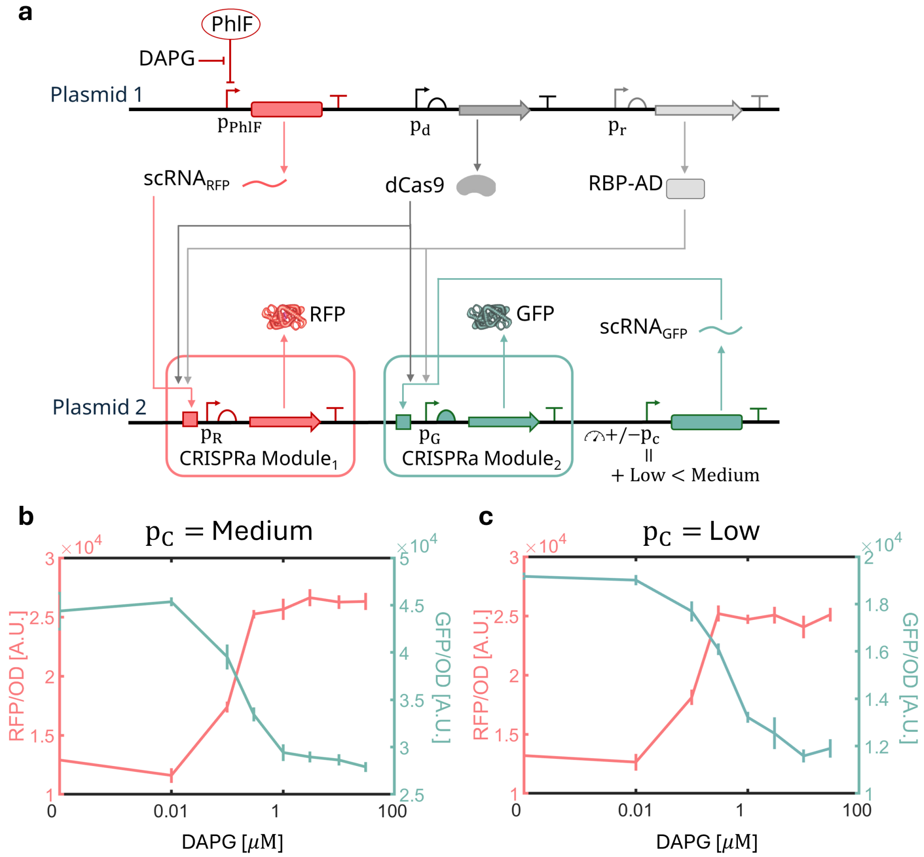


**SI Figure 9.** Coupling between two CRISPRa modules for medium and low competitor scRNA levels. (a) Genetic construct with two CRISPRa modules. CRISPRa Module 1 has input scRNA$_{RFP}$ and output RFP, while CRISPRa Module 2 has input scRNA$_{GFP}$ and output GFP. (b) Induction curve showing the level of output protein expressions (RFP in red and GFP in green) for medium levels of scRNA_GFP as we vary DAPG and therefore scRNA$_{RFP}$. (c) Induction curve showing the coupling between the modules for low scRNA$_{GFP}$. We see that the coupling persists for lower levels of the competitor scRNA.

#### 2.7 Fold change in RFP for different levels of DAPG

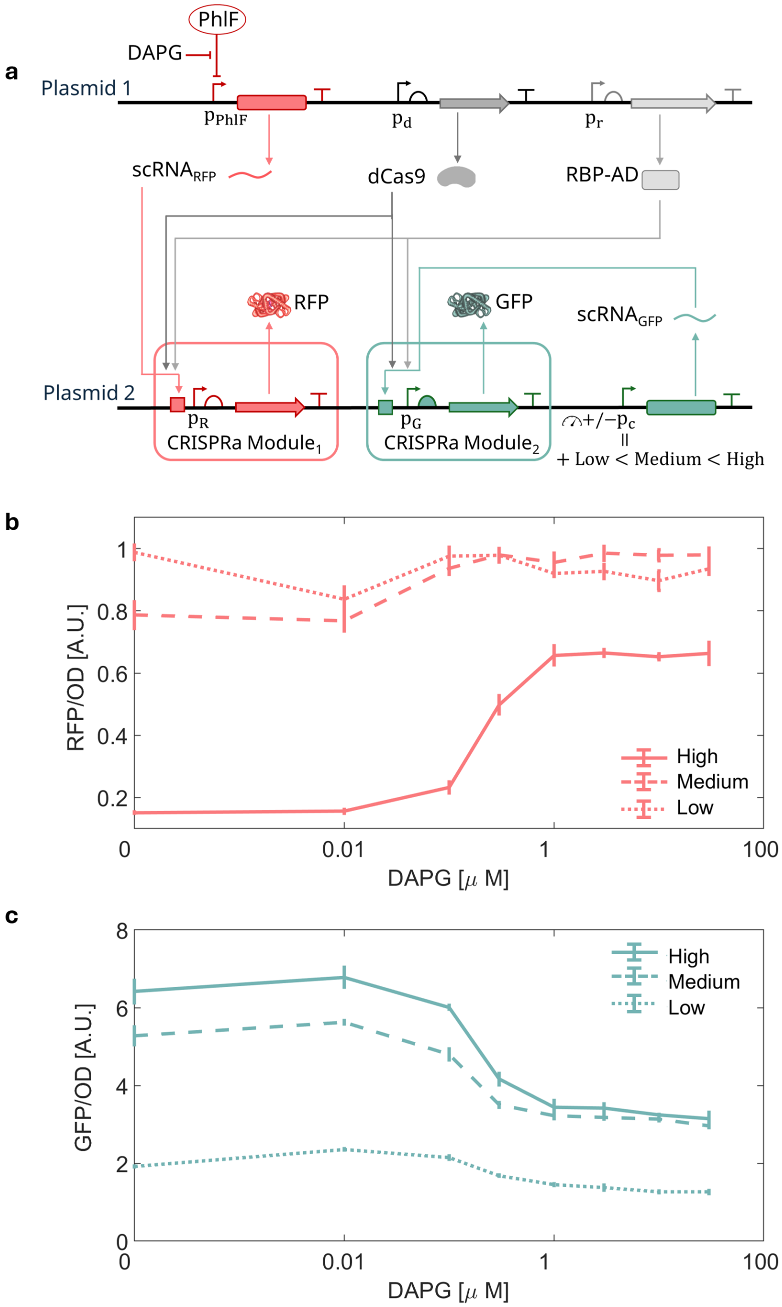


**SI Figure 10.** Drop in RFP due to competition for different levels of DAPG with low, medium and high levels of competitors $scRNA_{GFP}$ in the system. (a) Genetic construct with two CRISPRa modules. CRISPRa Module 1 has input $scRNA_{RFP}$ and output RFP, while CRISPRa Module 2 has input $scRNA_{GFP}$ and output GFP. The promoter of $scRNA_{GFP}$ is varied as low, medium, and high. (b) Fold change in RFP with respect to $scRNA_{RFP}$ for different promoter strength for the competitor scRNA. (c) Fold change in GFP with respect to $scRNA_{RFP}$ for different promoter strength for the competitor scRNA. The charts are normalized using the protein levels obtained in the absence of $scRNA_{GFP}$.

#### 2.8 Ruling out ribosome competition during concurrent CRISPRa activation

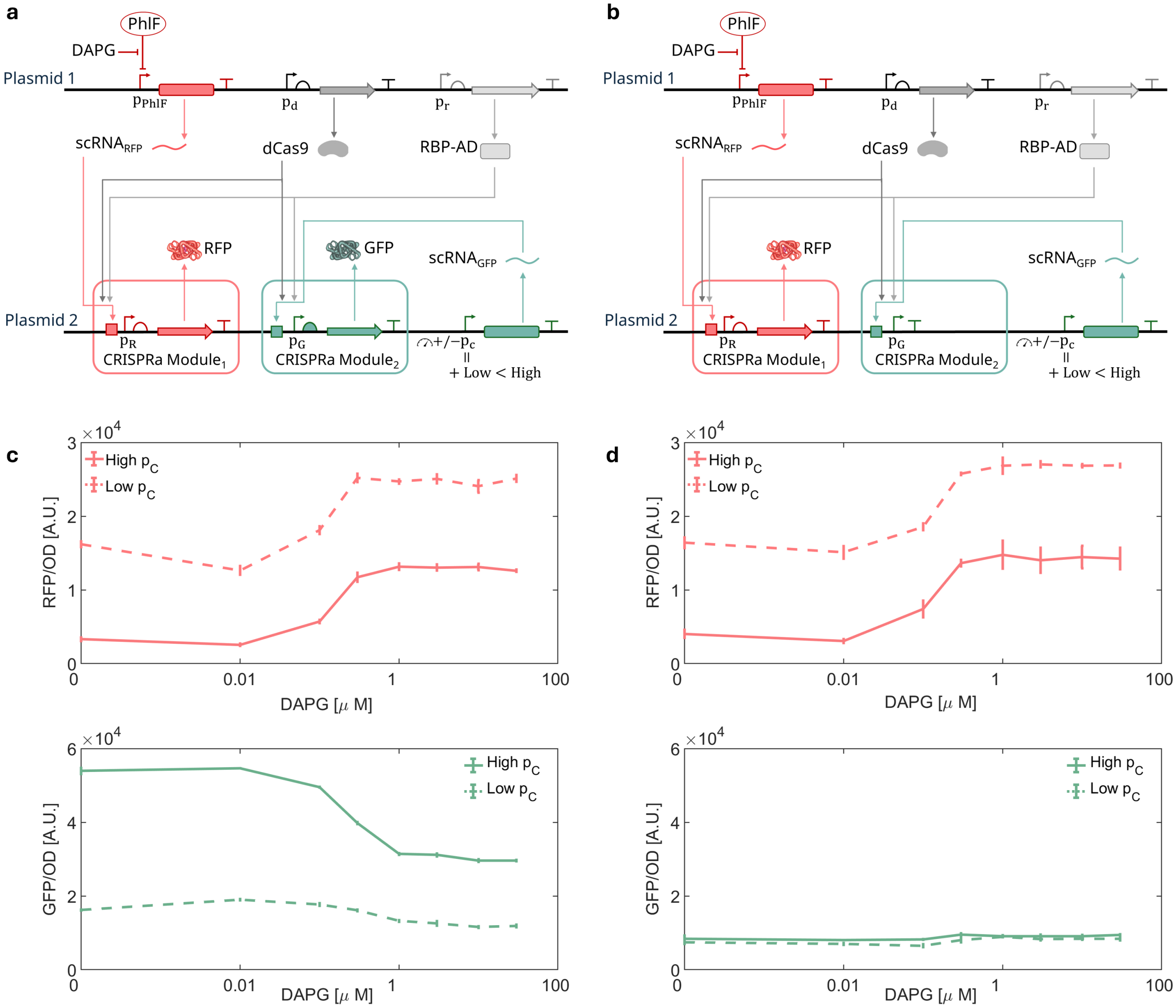


**SI Figure 11.** Coupling between two CRISPRa modules without ribosome competition between the modules. (a) Genetic construct with two CRISPRa modules, CRISPRa Module 1 has input $\mathrm{scRNA_{RFP}}$ and output RFP, while CRISPRa Module 2 has input $\mathrm{scRNA_{GFP}}$ and output GFP. (b) Genetic construct with two CRISPRa modules accounting for ribosome competition, CRISPRa Module 1 has input $\mathrm{scRNA_{RFP}}$ and output RFP, while CRISPRa Module 2 has input $\mathrm{scRNA_{GFP}}$ and no output protein expression. (c-Top) Input-output response of RFP for low and high levels of competitor $\mathrm{scRNA}_{GFP}$ with the expression of GFP. (c-Bottom) Off target response of GFP with DAPG for low and high levels of competitor $\mathrm{scRNA}_{GFP}$. We see the 85% drop in RFP at low DAPG and a 40% drop at high DAPG. (d-Top) Input-output response of RFP for low and high levels of competitor $\mathrm{scRNA}_{GFP}$ without the expression of GFP. (d-Bottom) Measured GFP response with DAPG for low and high levels of competitor $\mathrm{scRNA}_{GFP}$. The measured GFP values fluctuate around the background values verifying the absence of GFP in the system. Comparing (c) and (d), we see the 85% drop in RFP at low DAPG and a 40% drop at high DAPG is retained. Therefore, ribosome competition between GFP and RFP in construct (a) does not dictate the behavior of the CRISPRa system.

#### 2.9 Temporal data for each data point in Main Manuscript Figure 4

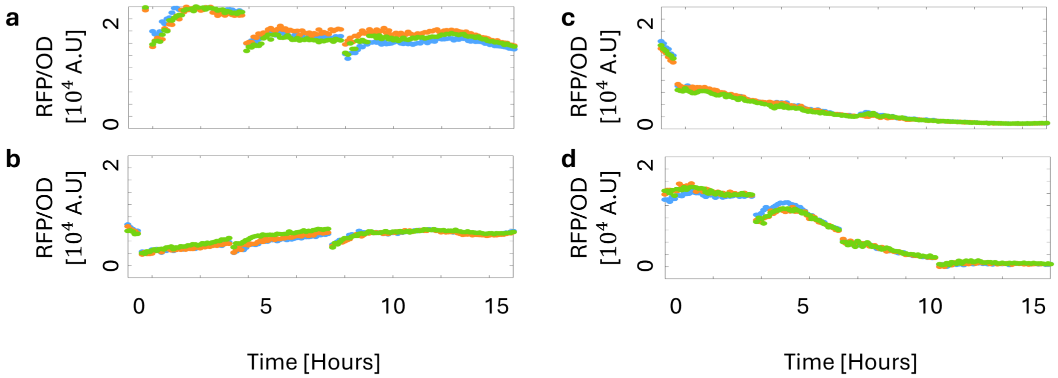


**SI Figure 12.** Time Series data for each data point in Main Manuscript Fig. 4(b) using the genetic construct shown in Main Manuscript Fig. 4(a). (a-d) Temporal evolution of RFP for the four different cases of scRNA$_{GFP}$ implementing selective resource competition. Here, the cases are as follows: in (a) scRNA$_{GFP}$ does not bind to dCas9 or RBP-AD, in (b) scRNA$_{GFP}$ binds to dCas9 only, in (c) scRNA$_{GFP}$ binds to RBP-AD only, and in (d) scRNA$_{GFP}$ binds to both dCas9 and RBP-AD. The steady states used in the Main Manuscript correspond to the time point when the measured OD reaches 0.14 in the final fourth batch.

#### 2.10 RFP and GFP responses for selective resource demand

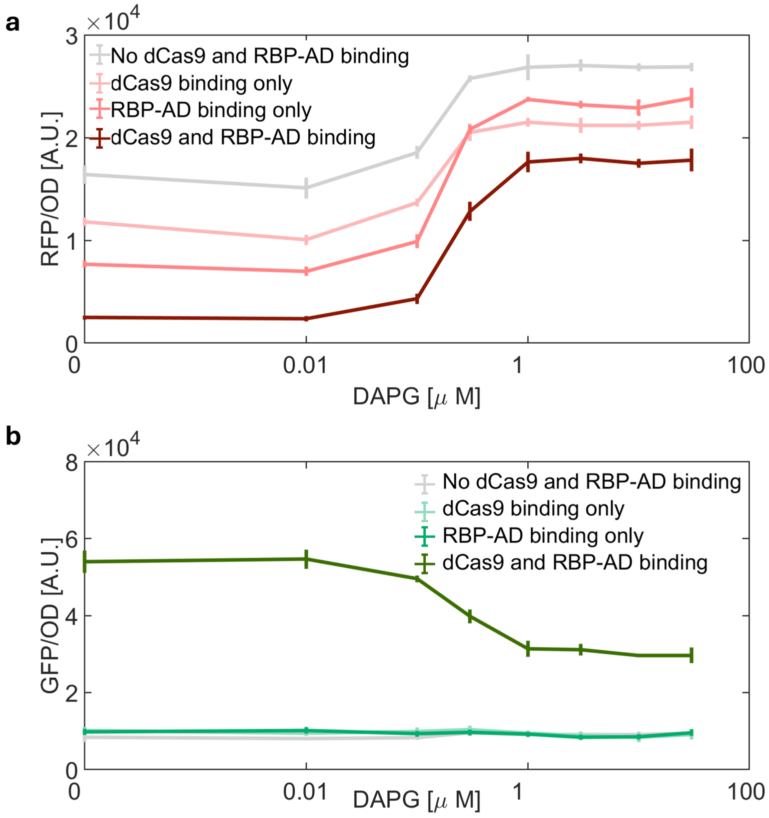


**SI Figure 13.** (a) On target response of RFP with increasing DAPG for four different variations of the scRNA$_{GFP}$. The variations are as follows: in case (I) scRNA$_{GFP}$ binds to dCas9 and RBP-AD, in case (II) scRNA$_{GFP}$ binds to RBP-AD only, in case (III) scRNA$_{GFP}$ binds to dCas9 only and in case (IV) scRNA$_{GFP}$ does not bind to dCas9 or RBP-AD. From the curves we see that case (I) with competition for both dCas9 and RBP-AD produces the least amount of RFP, whereas case (IV) without binding for both dCas9 and RBP-AD produces the most amount of RFP. Cases (II) and (III), with competition for one of the two resources stay in between the aforementioned extremities for all values of DAPG. (b) GFP levels measured for all the cases as DAPG is increased. GFP is produced only for case (I) and the levels decrease due to competition. GFP levels are near background values for Cases (II), (III), and (IV).

##### 2.11 Numerical simulations with selective resource demand

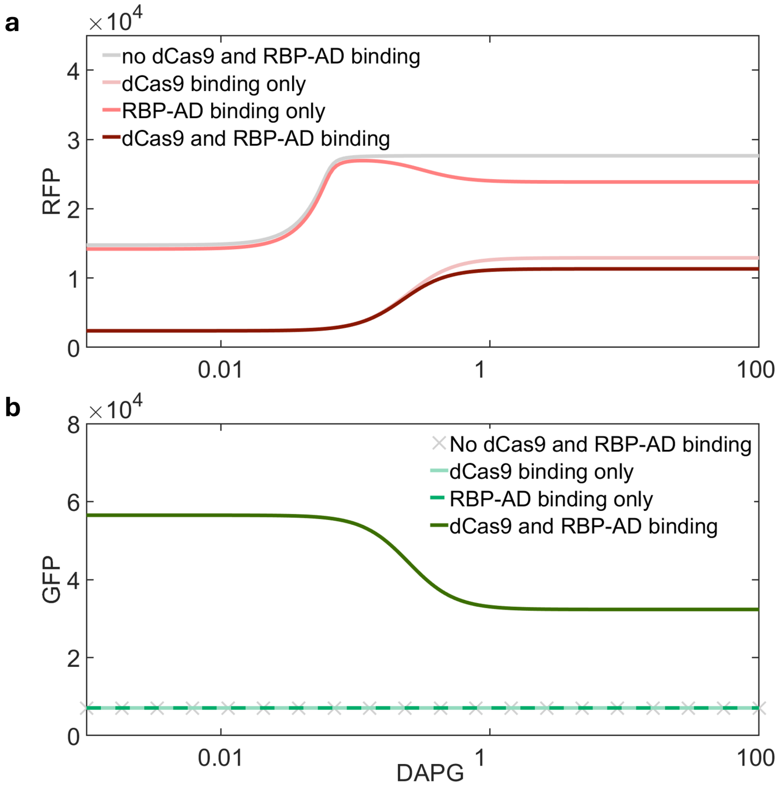


**SI Figure 14.** (a) Numerical simulations for the on-target response of RFP with increasing DAPG for four different variations of the scRNA$_{GFP}$. The variations are as follows: in case (I) scRNA$_{GFP}$ binds to dCas9 and RBP-AD, in case (II) scRNA$_{GFP}$ binds to RBP-AD only, in case (III) scRNA$_{GFP}$ binds to dCas9 only and in case (IV) scRNA$_{GFP}$ does not bind to dCas9 or RBP-AD. Case (II) and (III) are implemented by setting the forward binding reaction rates of the scRNA$_{GFP}$ with the corresponding resource as zero. From the curves we see that case (I) with competition for both dCas9 and RBP-AD produces the least amount of RFP, whereas case (IV) without binding for both dCas9 and RBP-AD produces the most amount of RFP. Cases (II) and (III), with competition for one of the two resources stay in between the aforementioned extremities for all values of DAPG. (b) Numerical simulations showing the GFP levels for all the cases as DAPG is increased. GFP is produced only for case (I) and the levels decrease due to competition. GFP levels are at the basal expression levels for Cases (II), (III), and (IV).

#### 2.12 dCas9 promoter characterization for Main Manuscript Figure 5

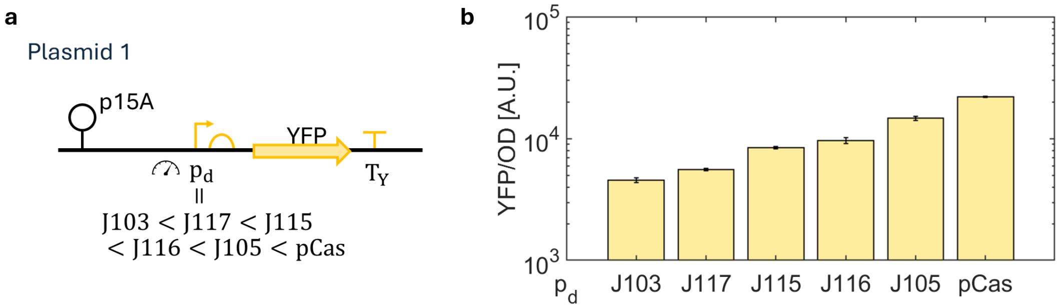


**SI Figure 15.** Promoter strength characterization for the promoters for dCas9 used in Main Manuscript Fig. 5. (a) Genetic construct producing YFP with the promoters to quantify the promoter strengths of dCas9 in Main Manuscript Fig. 5(a). The specific promoters are J103 (BBa_J23103), J117 (BBa_J23117), J115 (BBa_J23115), J116 (BBa_J23116), J105 (BBa_J23105), and pCas, see Table 5 for sequences. (b) Bar chart representation of YFP showing the relative strengths of the promoters used.

##### 2.13 Induction curve for different levels of dCas9

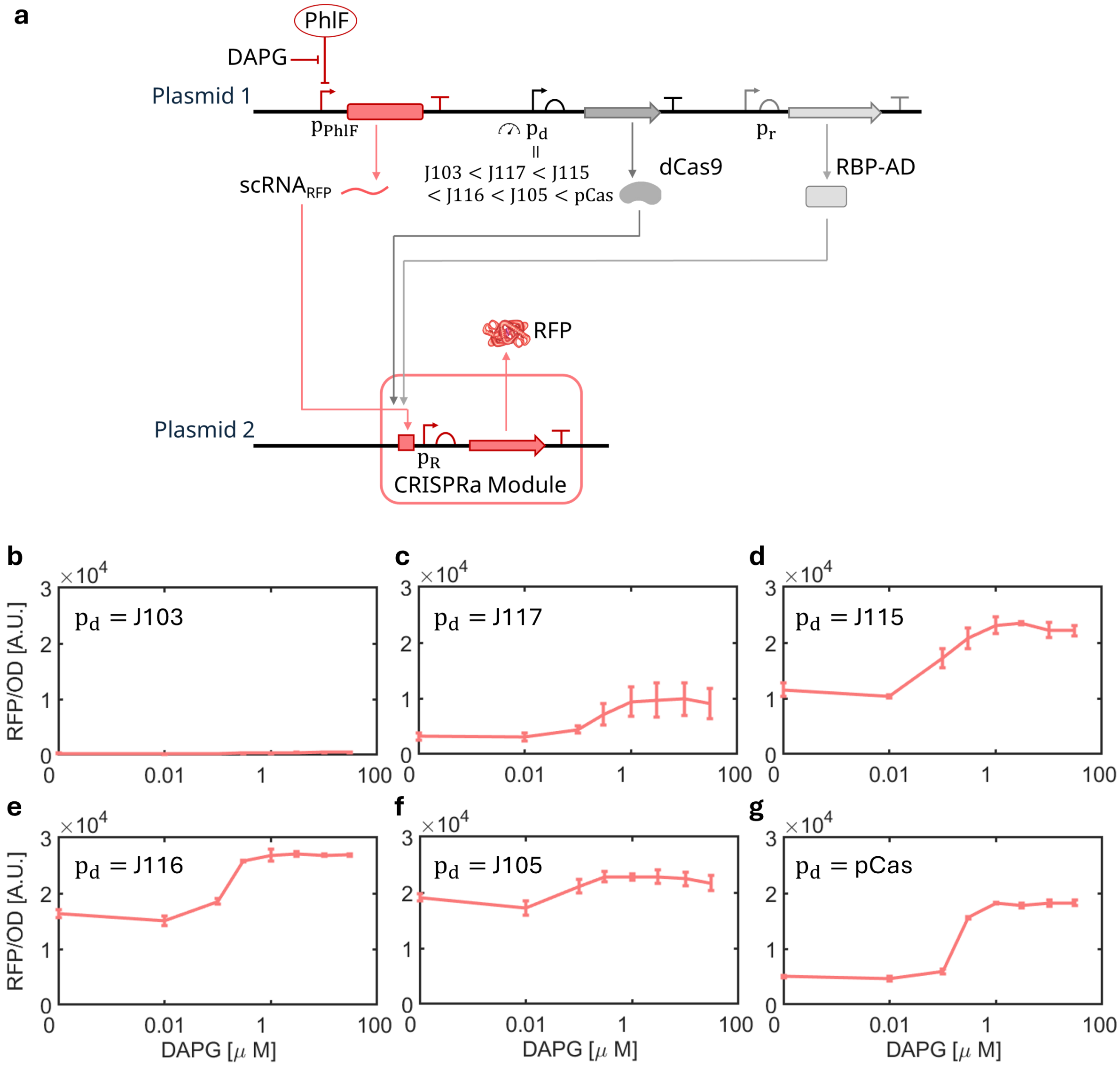


**SI Figure 16.** (a) Genetic construct with a single CRISPR module with inducibly produced scRNA$_{RFP}$ as input and RFP as the output for varying levels of dCas9. (b-g) Induction curve showing RFP with increasing DAPG for different promoters for dCas9 as: (b) $p_d$ = J103, (c) $p_d$ = J117, (d) $p_d$ = J115, (e) $p_d$ = J116, (f) $p_d$ = J105, and (g) $p_d$ = pCas.

#### 2.14 Numerical simulations of biphasic response with dCas9

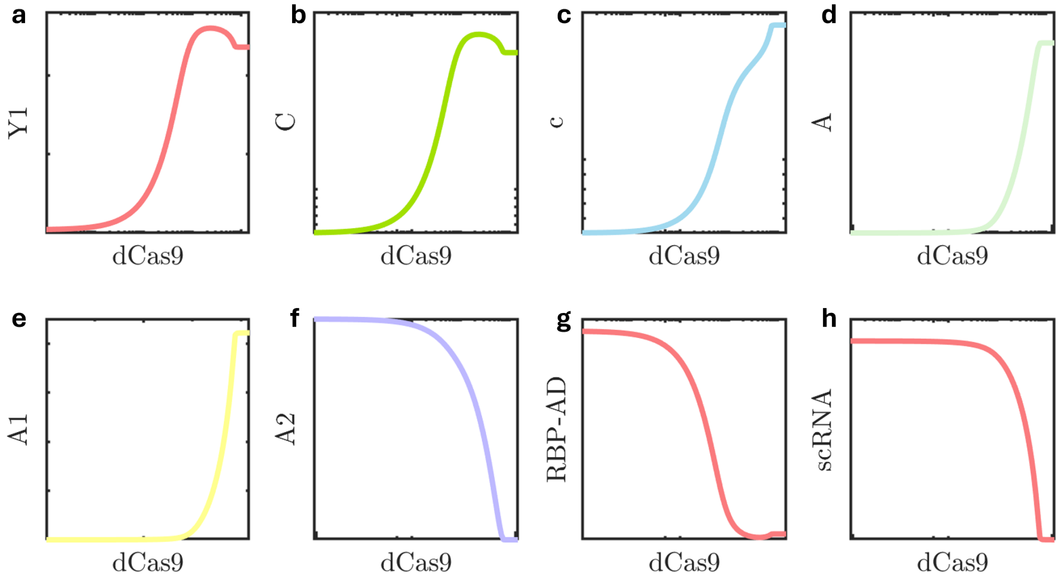


**SI Figure 17.** Line plots of levels of each complex and resources in a CRISPRa module as dCas9 is increased. The line plots correspond to (a) output protein ($Y$), (b) transcriptionally active complex ($C$), (c) intermediate scRNA-dCas9-DNA complex ($c$), (d) activator complex ($A$), (e) intermediate scRNA-dCas9 complex ($A_1$), (f) intermediate scRNA-RBP-AD complex ($A_2$), (g) free RBP-AD ($r$), and (h) free scRNA ($s$). From these plots, we see that as dCas9 is increased, the RBP-AD and the scRNA are being sequestered which leads to the biphasic response in $Y$. Specifically, as we increase the amounts of dCas9 the levels of intermediate complexes increase ($A$, $A_1$ and $c$) while free scRNA and RBP-AD decrease.

##### 2.15 Fold change in RFP and GFP for low and high dCas9

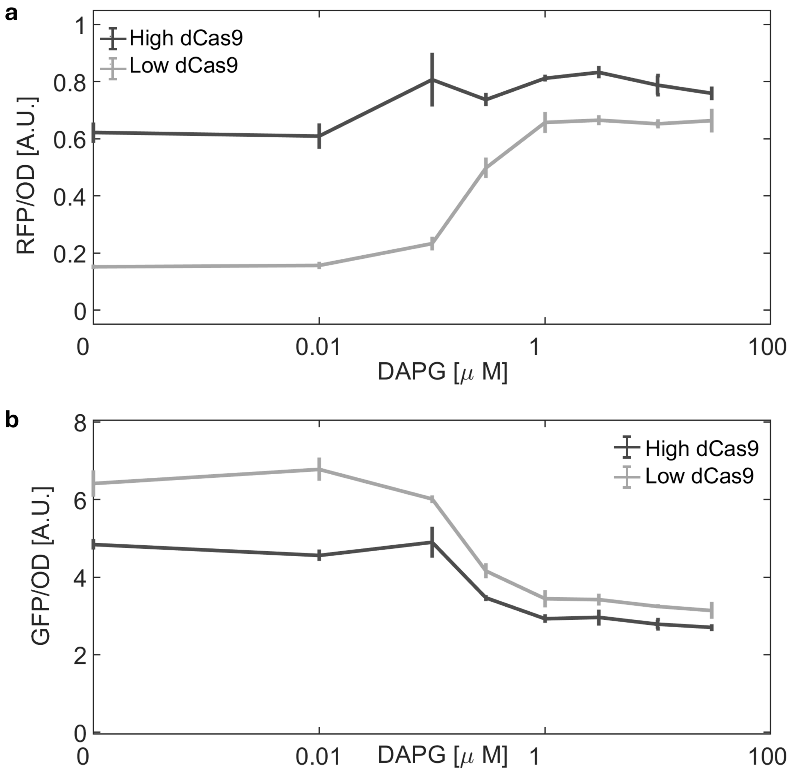


**SI Figure 18.** (a) Fold change in RFP for low and high dCas9 levels as DAPG (and thereby $scRNA_{RFP}$) is varied. The drop in RFP is higher for low dCas9 when compared to high dCas9. (b) Fold activation of GFP for low and high dCas9 levels as DAPG (and thereby $scRNA_{RFP}$) is varied. We observe that the fold activation drops significantly for both cases as DAPG is increased. Therefore, for all levels of DAPG, increasing dCas9 was able to reduce but not remove the drop in protein levels due to competition.

#### 2.16 Toxicity for dCas9 and RBP-AD

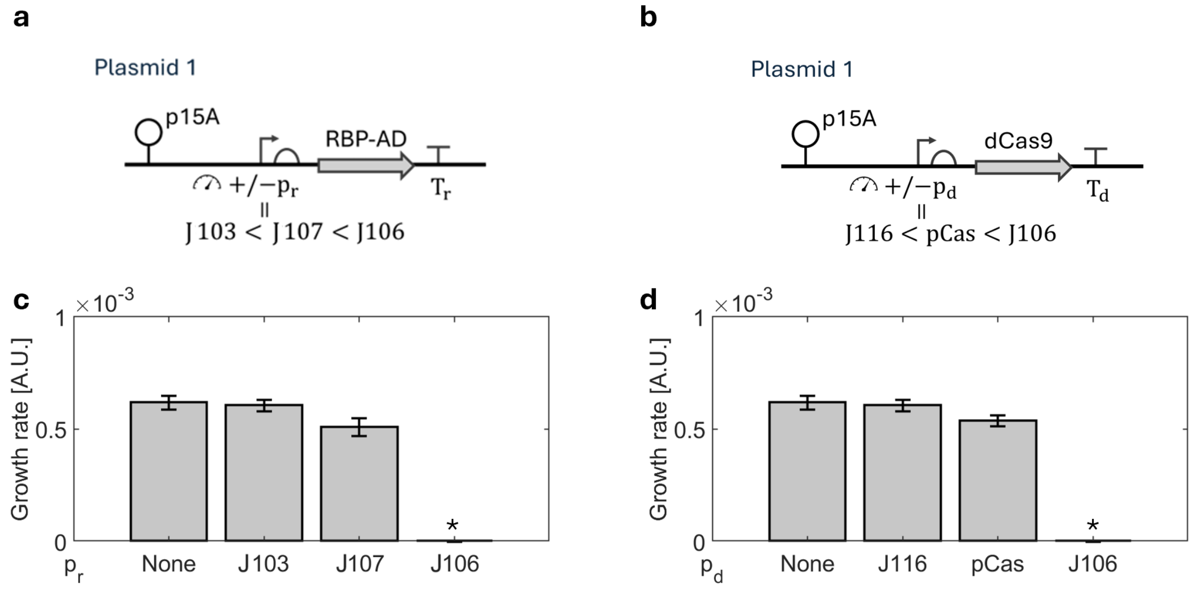


**SI Figure 19.** (a) Genetic construct with varying promoters for RBP-AD to check for RBP-AD toxicity. The promoters used are J103 (BBa_J23103), J107 (BBa_J23107), and J106 (BBa_J23106). (b) Genetic construct with varying promoters for dCas9 to check for dCas9 toxicity. The promoters used are J116 (BBa_J23116), pCas, and J106 (BBa_J23106). (c) Growth rate for varying levels of RBP-AD using the genetic construct in (a). The growth rate is calculated as $\frac{OD_{end}-OD_{start}}{time_{end}-timestart}$, where $OD_{end} = 0.12$, $OD_{start} = 0.02$. $time_{end}$ and $time_{start}$ are the time in minutes at which the background subtracted OD reached $OD_{end}$ and $OD_{start}$, respectively. (d) Growth rate for varying levels of dCas9 using the genetic construct in (b). The (*) on top of J106 in (c) and (d) indicates the lack of growth in the overnight cultures.

#### 2.17 Temporal data for each data point in Main Manuscript Figure 6(d)

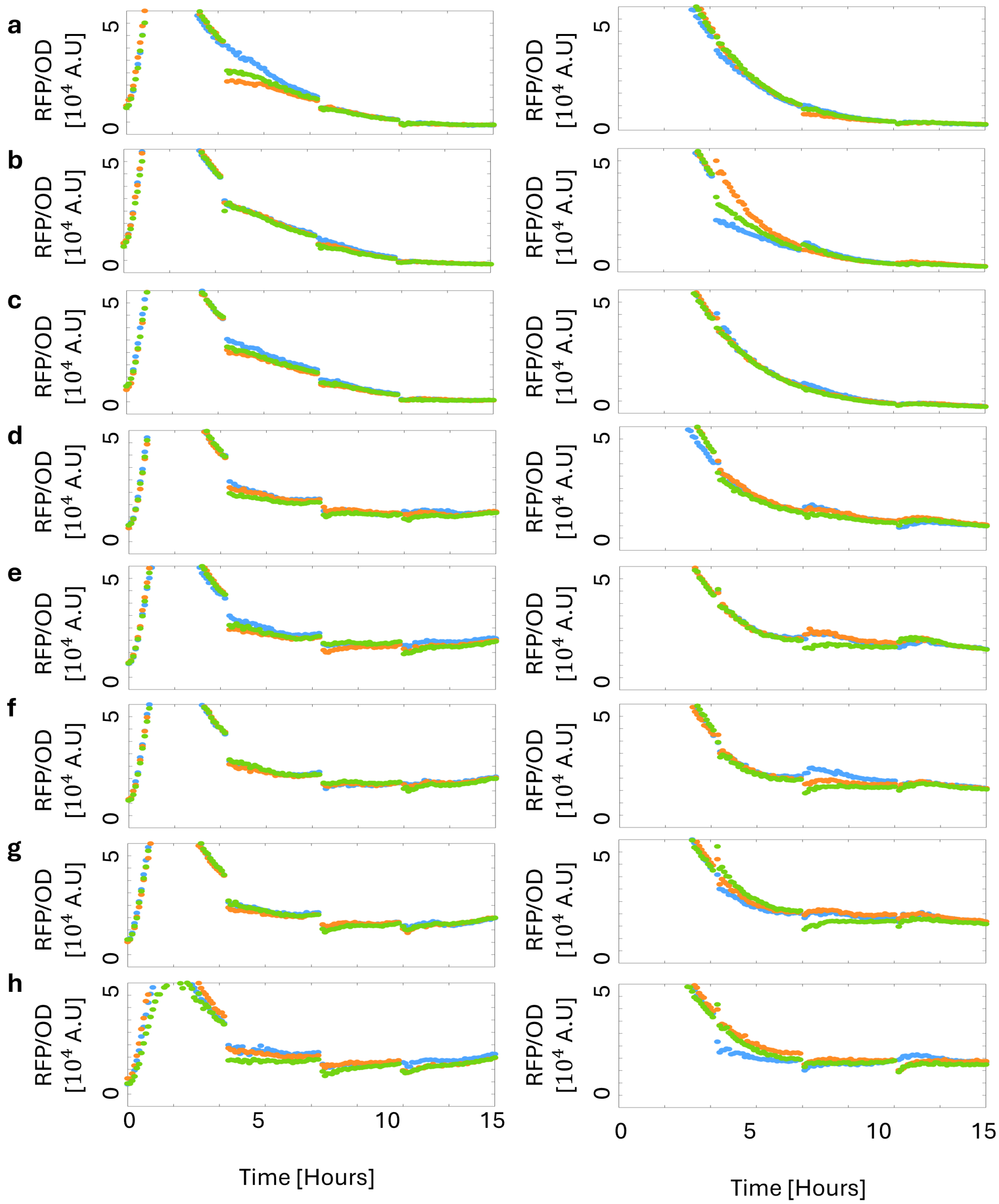


**SI Figure 20.** Time Series data for each data point in Main Manuscript Fig. 6(d) using the genetic construct shown in Main Manuscript Fig. 6(a). (a-h) Temporal evolution of RFP for the two different cases of scRNA$_{GFP}$ without a target site for different levels of DAPG. Here the target is designed such that (Left) has no PAM site and (Right) has no 20 bp complementary region to the scRNA binding site. The DAPG levels are (a) $0\mu M$, (b) $0.01\mu M$, (c) $0.1\mu M$, (d) $0.3\mu M$, (e) $1\mu M$, (f) $3\mu M$, (g) $10\mu M$, and (h) $30\mu M$, respectively. The steady states used in the Main Manuscript correspond to the time point when the measured OD reaches 0.14 in the final fourth batch.

#### 2.18 Reshaping the on-target response with competitor scRNA for high dCas9 level

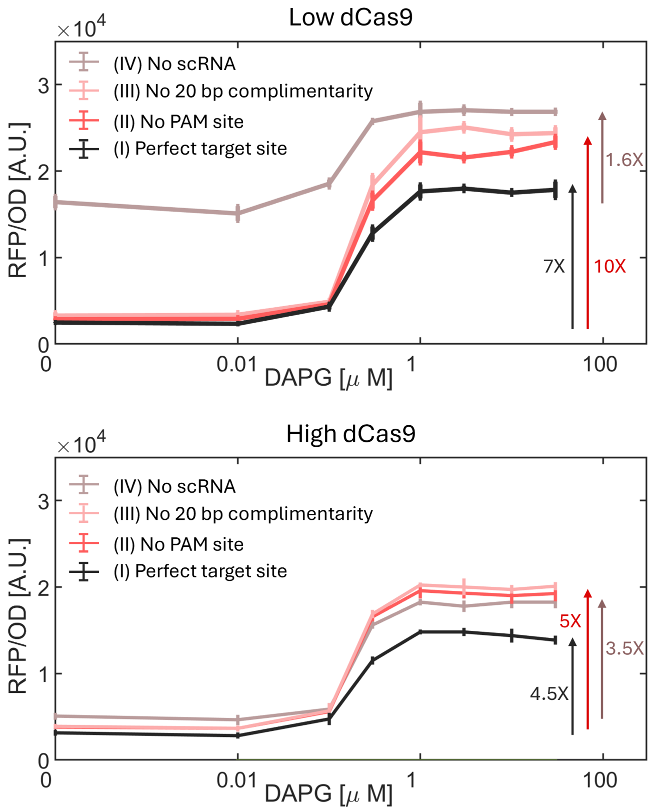


**SI Figure 21.** On target response of RFP with increasing DAPG using the construct shown in Main Manuscript Fig. 6(a) for (Top) low and (Bottom) high dCas9 levels. In the case of low dcas9, we see that the dynamic range of activation increased from 1.6-fold to 7-fold with the introduction of a competitor scRNA with a target site and further to 10-fold when the competitor scRNA does not have a target site. For the case of high dCas9, we see that the increase in fold change from no-competitor to having a competitor with a target site is 3.5-fold to 4.5-fold, while it increased further to 5-fold when using a competitor without the target site.

### 3 Supplementary Note 3: Mathematical model for CRISPRa

In this section, we establish a chemical reaction network-based model for CRISPRa genetic circuits[3]. Each CRISPRa module consists of an input scRNA ($s_i$) that regulates the production of the output protein, $Y_i$ (Fig. 22a). The resources, dCas9 and RBP-AD, are shared between the CRISPRa modules. Here, the scRNA binds either with the free dCas9 ($d$) in the system forming $A_{1,i}$ or the free RBP-AD resource ($r$) in the system forming $A_{2,i}$. Note, the scRNA must recruit dCas9 before binding with the target gene. The formed complex $A_{1,i}$, having recruited dCas9, may bind with the target gene ($D_i$) to form $c_i$, or with RBP-AD to form the CRISPR complex $A_i$. This is followed by the binding with RBP-AD and the target gene, respectively, to form the transcriptionally active complex $C_i$. On the other hand, the complex $A_{2,i}$ must combine with dCas9 forming $A_i$ followed by the combination with the target gene to form $C_i$. The transcriptionally active complex will then undergo transcription and translation producing the output protein $Y_i$. Therefore, the pathways from the input $s_i$ to its regulated output $Y_i$ can be portrayed as a double-diamond reaction network (Fig. 22b).

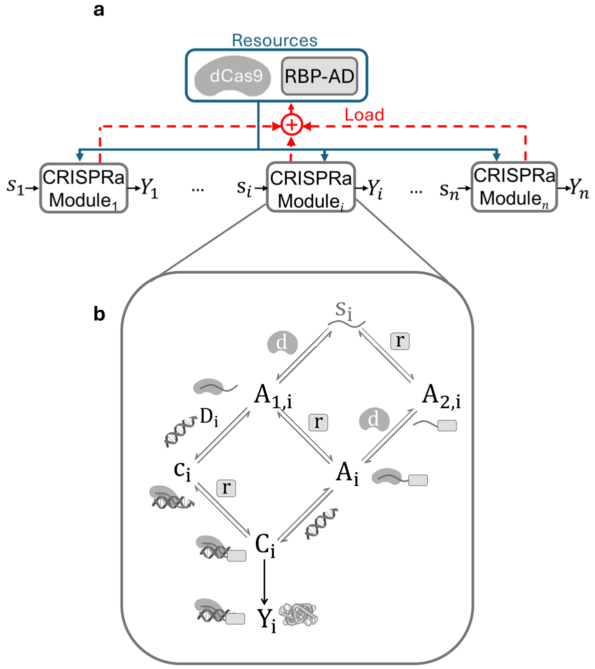


**SI Figure 22.** Double diamond reaction network with block diagram. (a) Block diagram representation of concurrent implementation of $n$ CRISPRa modules with scRNA $\{s_1, s_2, ..., s_n\}$ activating the production of proteins $\{Y_1, Y_2, ..., Y_n\}$ using the shared dCas9 and RBP-AD resources. The loads on the resource generator due to the consumption of the resources by each module are shown in red. (b) Chemical reaction diagram for modeling CRISPRa through physics-based reaction network models. It represents the pathways from the input scRNA ($s_i$) to the regulated output protein ($Y_i$) in a CRISPRa module and the complexes formed within each pathway. Double-sided arrows represent reversible binding reactions whereas the one-sided arrow from $C_i$ to $Y_i$ represents gene expression.

The reactions involved are as follows:

$$
\begin{aligned}
&\mathrm{d} + \mathrm{s}_i \underset{\mathrm{r}_i^-}{\overset{\mathrm{r}_i^+}{\rightleftharpoons}} \mathrm{A}_{1,i} && \mathrm{r} + \mathrm{s}_i \underset{\mathrm{p}_i^-}{\overset{\mathrm{p}_i^+}{\rightleftharpoons}} \mathrm{A}_{2,i} \\
&\mathrm{A}_{1,i} + \mathrm{D}_i \underset{\mathrm{t}_i^-}{\overset{\mathrm{t}_i^+}{\rightleftharpoons}} \mathrm{c}_i && \mathrm{A}_{1,i} + \mathrm{r} \underset{\mathrm{q}_i^-}{\overset{\mathrm{q}_i^+}{\rightleftharpoons}} \mathrm{A}_i \\
&\mathrm{A}_{2,i} + \mathrm{d} \underset{\mathrm{s}_i^-}{\overset{\mathrm{s}_i^+}{\rightleftharpoons}} \mathrm{A}_i && \mathrm{c}_i + \mathrm{r} \underset{\mathrm{h}_i^-}{\overset{\mathrm{h}_i^+}{\rightleftharpoons}} \mathrm{C}_i \\
&\mathrm{A}_i + \mathrm{D}_i \underset{\mathrm{g}_i^-}{\overset{\mathrm{g}_i^+}{\rightleftharpoons}} \mathrm{C}_i && \mathrm{C}_i \xrightarrow{\kappa} \mathrm{Y}_i + \mathrm{C}_i.
\end{aligned}
\tag{1}
$$

and the corresponding reaction rate equations can be obtained using mass action kinetics as:

$$
\begin{aligned}
\dot{Y}_i &= \kappa_i C_i + \beta_i D_i - \gamma Y_i \\
\dot{C}_i &= g_i^+ A_i D_i + h_i^+ c_i r - (h_i^- + g_i^- + \delta) C_i \\
\dot{c}_i &= t_i^+ A_{1,i} D_i + h_i^- C_i - (t_i^- + h_i^+ r + \delta) c_i \\
\dot{A}_i &= q_i^+ A_{1,i} r + s_i^+ A_{2,i} d + g_i^- C_i - (q_i^- + s_i^- + g_i^+ D_i + \delta) A_i \\
\dot{A}_{1,i} &= r_i^+ s_i d + q_i^- A_i + t_i^- c_i - (r_i^- + q_i^+ r + t_i^+ D_i + \delta) A_{1,i} \\
\dot{A}_{2,i} &= p_i^+ r s_i + s_i^- A_i - (p_i^- + s_i^+ d + \delta) A_{2,i} \\
\dot{s}_i &= u_i + r_i^- A_{1,i} + p_i^- A_{2,i} - (r_i^+ d + p_i^+ r + \delta) s_i.
\end{aligned} \tag{2}
$$

for $i \in \{1, 2, ..., n\}$, where, $\delta$ and $\gamma$ are the corresponding decay rate constants of scRNAs and proteins. Here, $\kappa$ is the activated production rate of the protein whereas $\beta$ is the basal production rate of the protein.

We define $r_t, d_t$ and $D_{it}$ as the total concentrations of RBP-AD, dCas9, and the target gene, respectively, their conservation in the system can be written as:

$$
\begin{aligned}
\text{DNA:} D_{it} &= D_i + C_i + c_i \\
\text{dCas9:} d_t &= d + \Sigma_i^n C_i + \Sigma_i^n A_i + \Sigma_i^n c_i + \Sigma_i^n A_{1,i} \\
\text{RBP-AD:} r_t &= r + \Sigma_i^n C_i + \Sigma_i^n A_i + \Sigma_i^n A_{2,i}.
\end{aligned}
$$

**Single CRISPRa module**

For a single CRISPRa module used in the numerical simulations in the Main Manuscript Fig. 1 and Fig. 2, we set $n = 1$ in equations 1 - 2. The conservation laws are modified as follows:

$$
\begin{aligned}
\text{DNA:} \quad & D_t = D + C + c \\
\text{dCas9:} \quad & d_t = d + C + A + c + A_1 \\
\text{RBP-AD:} \quad & r_t = r + C + A + A_2.
\end{aligned}
$$

**A pair of CRISPRa modules**

For a pair of CRISPRa modules used in the numerical simulations in the Main Manuscript Figs. 3 (c,d), we set $n = 2$ in equations 1 - 2, such that we have two sets of equations for $i = 1$ and $i = 2$. The conservation laws are modified as follows:

$$
\begin{aligned}
\text{DNA for module 1:} \quad & D_{1t} = D_1 + C_1 + c_1 \\
\text{DNA for module 2:} \quad & D_{2t} = D_2 + C_2 + c_2 \\
\text{dCas9:} \quad & d_t = d + C_1 + C_2 + A_1 + A_2 + c_1 + c_2 + A_{1,1} + A_{1,2} \\
\text{RBP-AD:} \quad & r_t = r + C_1 + C_2 + A_1 + A_2 + A_{2,1} + A_{2,2}.
\end{aligned}
$$

The production of scRNA$_{RFP}$ using the DAPG-inducible pPhlF promoter is modeled as:

$$
u_1(\text{DAPG}) = \alpha \frac{\text{DAPG}^{\text{N}}}{\text{K}^{\text{N}} + \text{DAPG}^{\text{N}}}.
$$

The implementation of no scRNA in Main Manuscript Fig. 3(e) is done by setting $u_2 = 0$. On the other hand, for the case of scrambled scRNA without dCas9 and RBP-AD binding, the forward reaction rate constants corresponding to the binding with dCas9 ($r_2^+, s_2^+$) and RBP-AD ($p_2^+, q_2^+, h_2^+$) are set to 0.

##### 3.1 Implementing selective resource binding of scRNA$_{GFP}$

To implement selective resource demand through modifications on scRNA$_{GFP}$ as shown in Main Manuscript Fig. 4(a) in the model, we fix the parameters for CRISPRa module 2 as follows.

| Case | Parameter varied |
|---|---|
| (I) scRNA$_{GFP}$ binding with dCas9 and RBP-AD | No changes |
| (II) scRNA$_{GFP}$ binding with RBP-AD only | $r_2^+ = s_2^+ = 0$ |
| (III) scRNA$_{GFP}$ binding with dCas9 only | $p_2^+ = q_2^+ = h_2^+ = 0$ |
| (IV) scrambled scRNA$_{GFP}$ (no dCas9 or RBP-AD binding) | $r_2^+ = s_2^+ = p_2^+ = q_2^+ = h_2^+ = 0$ |

##### 3.2 Modifying competitor scRNA to improve the response

To investigate different variations of the competitor scRNA for shaping the input/output response of a CRISPRa module, we compare the fold activation and the maximal activation produced by each scRNA variations. The competitor scRNA variations investigated along with the parameter fixed to obtain the change are as follows:

| Case | Parameter varied |
|---|---|
| No competitor scRNA | $u_2 = 0$ |
| With competitor scRNA | $u_2 = 7$ |
| Competitor scRNA without target site | $D_{2t} = 0$ |
| Competitor scRNA binding with dCas9 only | $p_2^+ = q_2^+ = h_2^+ = 0$ |
| Competitor scRNA binding with RBP-AD only | $r_2^+ = s_2^+ = 0$ |

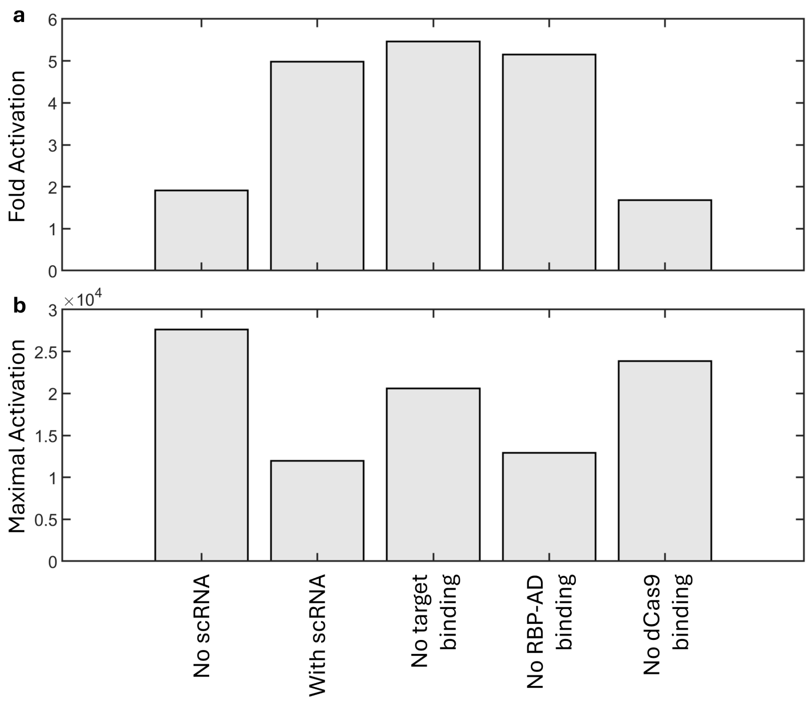


**SI Figure 23.** Model investigation of shaping the CRISPRa response. (a) Fold activation of CRISPRa module 1 obtained for different variations for the competitor scRNA. The variations investigated are as follows: no competitor scRNA, with competitor scRNA, competitor scRNA without a target, competitor scRNA binding with dCas9 only (no RBP-AD binding), and competitor scRNA binding with RBP-AD only (no dCas9 binding). (b) Maximal activation of CRISPRa module 1 obtained for different variations for the competitor scRNA. From (a) and (b), we can infer that adding a competitor scRNA without a target site produces the highest fold-change while maintaining high maximal activation levels.

##### 3.3 Model fitting to obtain parameters

To obtain the parameter values for the model, we performed model fitting using the fmincon function in MATLAB which provides the parameter set that minimizes a given function. The experimental data set used for fitting are:

- RFP and GFP data from Main Manuscript Fig. 3(c) (16 data points)
- RFP data from selective resource sharing from Fig. 15 (24 data points each)
- RFP data without scRNA binding site in module 2 from Main Manuscript Fig. 6(c,d) (8 data points)

The function to be minimized is the squared difference between the fitted model and the experimental data. The parameters fitted from the model are provided in Table 4.

Additionally, the values of scRNA (for Main Manuscript Fig. 1c), $p_C$ (for Main Manuscript Fig. 3d), and $d_t$ (for Main Manuscript Fig. 5) were chosen with appropriate scaling based on the promoter strength after obtaining the fitted parameters.

| Parameter | Symbol | Parameter Values | Reference |
|---|---|---|---|
| Total concentrations: | $D_t$ (DNA) | 7 for PSC101 and 80 for PSC101(E93G) | [2] |
| | $d_t$ (dCas9) | 1 | - |
| | $r_t$ (RBP-AD) | 30 | - |
| Activated Production Rate of RFP | $\kappa_1$ | 1000 | experiment |
| Basal Production Rate of RFP | $\beta_1$ | 100 | experiment |
| Activated Production Rate of GFP | $\kappa_2$ | 2000 | experiment |
| Basal Production Rate of GFP | $\beta_2$ | 350 | experiment |
| Dilution rate | $\gamma$ | $\frac{log(2)}{20}$ | [4] |
| scRNA decay rate | $\delta$ | 0.2 | [4] |
| DAPG Induction: | $\alpha$ | 5 | |
| | $K_m$ | 0.05 | [1] |
| | $N$ | 2.5 | |
| Reaction rate constants: | $p_i^+$ | 20 | - |
| | $p_i^-$ | 0.2 | - |
| | $r_i^+$ | 30 | - |
| | $r_i^-$ | 0.03 | - |
| | $t_i^+$ | 1 | - |
| | $t_i^-$ | 0.5 | - |
| | $h_i^+$ | 30 | - |
| | $h_i^-$ | 0.0003 | - |

**Table 4.** Parameters obtained through model fitting. Note: parameters such as total DNA (copy number), degradation rate, dilution rate, and the ones for DAPG induction are obtained from the appropriate references.

#### 4 Supplementary Note 4: DNA Sequences

| Promoter Name | Sequence |
|---|---|
| BBa_J23116 | GAATTCGCGGCCGCTTCTAGAGTTGACAGCTAGCTCAGTCCTAGGGACTATGCTAGC |
| BBa_J23103 | GGCGCGCCCTGATAGCTAGCTCAGTCC |
| BBa_J23117 | TTGACAGCTAGCTCAGTCCTAGGGATTGTGCTAGC |
| BBa_J23115 | TAGCTAGCTCAGCCCTTGGTACAATGCTAGC |
| BBa_J23105 | CGGCTAGCTCAGTCCTAGGTACTATGCTAGC |
| Sp.pCas9 | Provided later with dCas9 sequence |
| pTac | TTGACAATTAATCATCGGCTCGTATAATGTGTGGAATTGTGAGCGCTCACAATTAG |
| pPhlF | CGACGTACGGTGGAATCTGATTCGTTACCAATTGACATGATACGAAACGTACCGTATCGTTAAGGT |
| BBa_J23110 | TTTACGGCTAGCTCAGTCCTAGGTACAATGCTAGC |
| BBa_J23119 | TTGACAGCTAGCTCAGTCCTAGGTATAATACTAGT |
| BBa_J23107 | TTTACGGCTAGCTCAGCCCTAGGTATTATGCTAGC |
| BBa_J23114 | GGCGCGCCTTTATGGCTAGCTCAGTCCTAGGTACAATGCTAGCC |
| LacUV5 | GGCGCGCCCCCCAGGCTTTACACTTTATGCTTCCGGCTCGTATAATGTGTGG |
| pTrc | GGCGCGCCTTGACAATTAATCATCCGGCTCGTATAATGTGTGG |

**Table 5.** Promoter Sequences

| RBS Name | Sequence |
|---|---|
| BBa_B0034 | TACTAGAGAAAGAGGAGAAAGGATCT |
| BBa_J34801 | GAATTCATTAAAGAGGAGAAAGGTACC |
| RBS_RFP | GAATTCATTAAAGAGGAGAAAGGTACC |
| RBS_GFP | TCATACCGACTCGTAGGGGGTAATAA |

**Table 6.** RBS Sequences

| Terminator Name | Sequence |
|---|---|
| BBa_B0015 | CTCGAGTAAGGATCTCCAGGCATCAAATAAAACGAAAGGCTCAGTCGAAAGACTGGGCC TTTCGTTTTATCTGTTGTTTGTCGGTGAACGCTCTCTACTAGAGTCACACTGGCTCACCTT CGGGTGGGCCTTTCTGCG |
| BBa_B1002 | CGCAAAAAACCCCGCTTCGGCGGGGTTTTTTCGC |
| rrnB T1 | CAAATAAAACGAAAGGCTCAGTCGAAAGACTGGGCCTTTCGTTTTATCTGTTGTTTGTCG GTGAACGCTCTC |
| L3S3P12 | CCAATTATTGAACACCCGAAAGGGTGTTTTTTTGTTTCTGGTCTCCC |
| BBa_K1893035 | GAAGCTTGGGCCCGAACAAAAACTCATCTCAGAAGAGGATCTGAATAGCGCCGTCGACC ATCATCATCATCATCATTGAGTTTAAACGGTCTCCAGCTTGGCTGTTTTGGCGGATGAGAG AAGATTTTCAGCCTGATACAGATTAAATCAGAACGCAGAAGCGGTCTGATAAAACAGAAT TTGCCTGGCGGCAGTAGCGCGGTGGTCCCACCTGACCCCATGCCGAACTCAGAAGTGAA ACGCCGTAGCGCCGATGGTAGTGTGGGGTCTCCCCATGCGAGAGTAGGGAACTGCCAGG CATCAAATAAAACGAAAGGCTCAGTCGAAAGACTGGGCCTTTCGTTTTATCTGTTGTTTG TCGGTGAACT |
| T7Te | GGCTCACCTTCGGGTGGGCCTTTCTGCG |
| L3S2P21 | CTCGGTACCAAATTCCAGAAAAGAGGCCTCCCGAAAGGGGGGCCTTTTTCGTTTTGGTCC |

**Table 7.** Terminator Sequences

#### 4.1 Protein Sequences

**RFP reporter gene:**

The mRFP1 reporter gene was used in Main Manuscript Figs. 1,3,4,5,6. It contains a J306 binding sequence containing a PAM site allowing the $scRNA_{RFP}$ to bind to the target at -81 bp to the TSS on the non-template strand, which was reported to produce optimal activation[5].

PAM site + J306 binding sequence,BBa_J23117 promoter,RBS_RFP, mRFP1, rrnB T1 terminator

CCTACGGAGCGTTCTGGACACAACGTCGTCTTGAAGTTGCGATTATAGATTGACA
GCTAGCTCAGTCCTAGGGATTGTGCTAGCGAATTCATTAAAGAGGAGAAAGGTAC
CATGGCGAGTAGCGAAGACGTTATCAAAGAGTTCATGCGTTTCAAAGTTCGTATG
GAAGGTTCCGTTAACGGTCACGAGTTCGAAATCGAAGGTGAAGGTGAAGGTCGT
CCGTACGAAGGTACCCAGACCGCTAAACTGAAAGTTACCAAAGGTGGTCCGCTGC
CGTTCGCTTGGGACATCCTGTCCCCGCAGTTCCAGTACGGTTCCAAAGCTTACTT
AAACACCCGGCTGACATCCCGGACTACCTGAAACTGTCCTTCCCGGAAGGTTTCA
AATGGGAACGTGTTATGAACTTCGAAGACGGTGGTGTTGTTACCGTTACCCAGGA
CTCCTCCCTGCAAGACGGTGAGTTCATCTACAAAGTTAAACTGCGTGGTACCAAC
TTCCCGTCCGACGGTCCGGTTATGCAGAAAAAAACCATGGGTTGGGAAGCTTCCA
CCGAACGTATGTACCCGGAAGACGGTGCTCTGAAAGGTGAAATCAAAATGCGTCT
GAAACTGAAAGACGGTGGTCACTACGACGCTGAAGTTAAAACCACCTACATGGCT
AAAAAACCGGTTCAGCTGCCGGGTGCTTACAAAACCGACATCAAACTGGACATCA
CCTCCCACAACGAAGACTACACCATCGTTGAACAGTACGAACGTGCTGAAGGTCG
TCACTCCACCGGTGCTTAAGGATCCAAACTCGAGTAAGGATCTCCAGGCATCAAA
TAAAACGAAAGGCTCAGTCGAAAGACTGGGCCTTTCGTTTTATCTGTTGTTTGTC
GGTGAACGCTCTC

**GFP reporter gene**:

The ffGFP reporter gene was used in Main Manuscript Figs. 3,4,5,6. It contains a J108 binding sequence containing a PAM site allowing the $scRNA_{GFP}$ to bind to the target. Modifications that are made to this binding region are specified in the following table.

PAM site + J108 binding sequence,BBa_J23117 promoter,RBS_GFP, ffGFP, L3S2P21 terminator

CCTGCGGTGTCCTGCGGTTACCACGTCGTCTTGAAGTTGCGATTATAGATTGACAG
CTAGCTCAGTCCTAGGGATTGTGCTAGCTCATACCGACTCGTAGGGGGTAATAAA
TGCGTAAAGGCGAAGAGCTGTTCACTGGTGTCGTCCCTATTCTGGTGGAACTGGA
TGGTGATGTCAACGGTCATAAGTTTTCCGTGCGTGGCGAGGGTGAAGGTGACGCA
ACTAATGGTAAACTGACGCTGAAGTTCATCTGTACTACTGGTAAACTGCCGGTAC
CTTGGCCGACTCTGGTAACGACGCTGACTTATGGTGTTCAGTGCTTTGCTCGTTAT
CCGGACCATATGAAGCAGCATGACTTCTTCAAGTCCGCCATGCCGGAAGGCTATG
TGCAGGAACGCACGATTTCCTTTAAGGATGACGGCACGTACAAAACGCGTGCGGA
AGTGAAATTTGAAGGCGATACCCTGGTAAACCGCATTGAGCTGAAAGGCATTGAC
TTTAAAGAAGACGGCAATATCCTGGGCCATAAGCTGGAATACAATTTTAACAGCC
ACAATGTTTACATCACCGCCGATAAACAAAAAAATGGCATTAAAGCGAATTTTAA
AATTCGCCACAACGTGGAGGATGGCAGCGTGCAGCTGGCTGATCACTACCAGCAA
AACACTCCAATCGGTGATGGTCCTGTTCTGCTGCCAGACAATCACTATCTGAGCA
CGCAAAGCGTTCTGTCTAAAGATCCGAACGAGAAACGCGATCATATGGTTCTGCT
GGAGTTCGTAACCGCAGCGGGCATCACGCATGGTATGGATGAACTGTACAAATAA
TAACTCGGTACCAAATTCCAGAAAAGAGGCCTCCCGAAAGGGGGGCCTTTTTTCGT
TTTGGTCC

For the Main Manuscript Fig. 6, modifications are made to the $scRNA_{GFP}$ binding site to implement 4 conditions. The sequences corresponding to each are provided here.

PAM Site + 20 bp complementary region

| Variation Name | Sequence |
|---|---|
| (I) Perfect target site for GFP | CCT GCGGTGTCCTGCGGTTACCA |
| (II) No PAM site for GFP | TTT GCGGTGTCCTGCGGTTACCA |
| (III) No 20 bp complementarity for GFP | CCT GAGGACGTGTTCGGCTACTA |
| (IV) No scRNA | TTT GAGGACGTGTTCGGCTACTA |

**Table 8.** $scRNA_{GFP}$ binding site variations in Fig. 6

**dCas9 gene:**

The dCas9 gene was used in all the figures in Main Manuscript. S. pyogenes dCas9 was expressed from its endogenous S. pyogenes Cas9 promoter or other promoters listed in Table 2.

S. pyogenes Cas9 promoter + RBS, dCas9, BBa_B0015 terminator

ACGTCTCATTTTCGCCAGATATCGACGTCTTAAGTTACGAAATCATCCTGTGGAG
CTTAGTAGGTTTAGCAAGATGGCAGCGCCTAAATGTAGAATGATAAAAGGATTAA
GAGATTAATTTCCCTAAAAATGATAAAACAAGCGTTTTGAAAGCGCTTGTTTTTT
TGGTTTGCAGTCAGAGTAGAATAGAAGTATCAAAAAAAGCACCGACTCGGTGCC
ACTTTTTCAAGTTGATAACGGACTAGCCTTATTTTAACTTGCTATGCTGTTTTGAA
TGGTTCCAACAAGATTATTTTATAACTTTTATAACAAATAATCAAGGAGAAATTC
AAAGAAATTTATCAGCCATAAAACAATACTTAATACTATAGAATGATAACAAAAT
AAACTACTTTTTAAAAGAATTTTGTGTTATAATCTATTTATTATTAAGTATTGGGT
AATATTTTTTGAAGAGATATTTTGAAAAAGAAAAATTAAAGCATATTAAACTAAT
TTCGGAGGTCATTAAAACTATTATTGAAATCATCAAACTCATTATGGATTTAATTT
AAACTTTTTATTTTAGGAGGCAAAAATGGATAAGAAATACTCAATAGGCTTAGCT
ATCGGCACAAATAGCGTCGGATGGGCGGTGATCACTGATGAATATAAGGTTCCGT
CTAAAAAGTTCAAGGTTCTGGGAAATACAGACCGCCACAGTATCAAAAAAAATCT
TATAGGGGCTCTTTTATTTGACAGTGGAGAGACAGCGGAAGCGACTCGTCTCAAA
CGGACAGCTCGTAGAAGGTATACACGTCGGAAGAATCGTATTTGTTATCTACAGG
AGATTTTTTCAAATGAGATGGCGAAAGTAGATGATAGTTTCTTTCATCGACTTGA
AGAGTCTTTTTTGGTGGAAGAAGACAAGAAGCATGAACGTCATCCTATTTTTGGA
AATATAGTAGATGAAGTTGCTTATCATGAGAAATATCCAACTATCTATCATCTGC
GAAAAAAATTGGTAGATTCTACTGATAAAGCGGATTTGCGCTTAATCTATTTGGC
CTTAGCGCATATGATTAAGTTTCGTGGTCATTTTTTGATTGAGGGAGATTTAAATC
CTGATAATAGTGATGTGGACAAACTATTTATCCAGTTGGTACAAACCTACAATCA
ATTATTTGAAGAAAACCCTATTAACGCAAGTGGAGTAGATGCTAAAGCGATTCTT
TCTGCACGATTGAGTAAATCAAGACGATTAGAAAATCTCATTGCTCAGCTCCCCG
GTGAGAAGAAAAATGGCTTATTTGGGAATCTCATTGCTTTGTCATTGGGTTTGAC
CCCTAATTTTAAATCAAATTTTGATTTGGCAGAAGATGCTAAATTACAGCTTTCA
AAAGATACTTACGATGATGATTTAGATAATTTATTGGCGCAAATTGGAGATCAAT
ATGCTGATTTGTTTTTGGCAGCTAAGAATTTATCAGATGCTATTTTACTTTCAGAT
ATCCTAAGAGTAAATACTGAAATAACTAAGGCTCCCCTATCAGCTTCAATGATTA
AACGCTACGATGAACATCATCAAGACTTGACTCTTTTAAAAGCTTTAGTTCGACA
ACAACTTCCAGAAAAGTATAAAGAAATCTTTTTTGATCAATCAAAAAACGGATAT
GCAGGTTATATTGATGGGGGAGCTAGCCAAGAAGAATTTTATAAATTTATCAAAC
CAATTTTAGAAAAAATGGATGGTACTGAGGAATTATTGGTGAAACTAAATCGTGA
AGATTTGCTGCGCAAGCAACGGACCTTTGACAACGGCTCTATTCCCCATCAAATT
CACTTGGGTGAGCTGCATGCTATTTTGAGAAGACAAGAAGACTTTTATCCATTTT
TAAAAGACAATCGTGAGAAGATTGAAAAAATCTTGACTTTTCGAATTCCTTATTA
TGTTGGTCCATTGGCGCGTGGCAATAGTCGTTTTGCATGGATGACTCGGAAGTCT
GAAGAAACAATTACCCCATGGAATTTTGAAGAAGTTGTCGATAAAGGTGCTTCAG
CTCAATCATTTATTGAACGCATGACAAACTTTGATAAAAATCTTCCAAATGAAAA
AGTACTACCAAAACATAGTTTGCTTTATGAGTATTTTACGGTTTATAACGAATTG
ACAAAGGTCAAATATGTTACTGAAGGAATGCGAAAACCAGCATTTCTTTCAGGTG
AACAGAAGAAAGCCATTGTTGATTTACTCTTCAAAACAAATCGAAAAGTAACCGT
TAAGCAATTAAAAGAAGATTATTTCAAAAAAATAGAATGTTTTGATAGTGTTGAA
ATTTCAGGAGTTGAAGATAGATTTAATGCTTCATTAGGTACCTACCATGATTTGCT
AAAAATTATTAAAGATAAAGATTTTTTGGATAATGAAGAAAATGAAGATATCTTA
GAGGATATTGTTTTAACATTGACCTTATTTGAAGATAGGGAGATGATTGAGGAAA
GACTTAAAACATATGCTCACCTCTTTGATGATAAGGTGATGAAACAGCTTAAACG
TCGCCGTTATACTGGTTGGGGACGTTTGTCTCGAAAATTGATTAATGGTATTAGG
GATAAGCAATCTGGCAAAACAATATTAGATTTTTTGAAATCAGATGGTTTTGCCA
ATCGCAATTTTATGCAGCTGATCCATGATGATAGTTTGACATTTAAAGAAGACAT
TCAAAAAGCACAAGTGTCTGGACAAGGCGATAGTTTACATGAACATATTGCAAAT

TTAGCTGGTAGCCCTGCTATTAAAAAAGGTATTTTACAGACTGTAAAAGTTGTTG
ATGAATTGGTCAAAGTAATGGGGCGGCATAAGCCAGAAAATATCGTTATTGAAAT
GGCACGTGAAAATCAGACAACTCAAAAGGGCCAGAAAAATTCGCGAGAGCGTAT
GAAACGAATCGAAGAAGGTATCAAAGAATTAGGAAGTCAGATTCTTAAAGAGCA
TCCTGTTGAAAATACTCAATTGCAAAATGAAAAGCTCTATCTCTATTATCTCCAA
AATGGAAGAGACATGTATGTGGACCAAGAATTAGATATTAATCGTTTAAGTGATT
ATGATGTCGATGCCATTGTTCCACAAAGTTTCCTTAAAGACGATTCAATAGACAA
TAAGGTCTTAACGCGTTCTGATAAAAATCGTGGTAAATCGGATAACGTTCCAAGT
GAAGAAGTAGTCAAAAAGATGAAAAACTATTGGAGACAACTTCTAAACGCCAAG
TTAATCACTCAACGTAAGTTTGATAATTTAACGAAAGCTGAACGTGGAGGTTTGA
GTGAACTTGATAAAGCTGGTTTTATCAAACGCCAATTGGTTGAAACTCGCCAAAT
CACTAAGCATGTGGCACAAATTTTGGATAGTCGCATGAATACTAAATACGATGAA
AATGATAAACTTATTCGAGAGGTTAAAGTGATTACCTTAAAATCTAAATTAGTTT
CTGACTTCCGAAAAGATTTCCAATTCTATAAAGTACGTGAGATTAACAATTACCA
TCATGCCCATGATGCGTATCTAAATGCCGTCGTTGGAACTGCTTTGATTAAGAAA
TATCCAAAACTTGAATCGGAGTTTGTCTATGGTGATTATAAAGTTTATGATGTTC
GTAAAATGATTGCTAAGTCTGAGCAAGAAATAGGCAAAGCAACCGCAAAATATT
TCTTTTACTCTAATATCATGAACTTCTTCAAAACAGAAATTACACTTGCAAATGG
AGAGATTCGCAAACGCCCTCTAATCGAAACTAATGGGGAAACTGGAGAAATTGTC
TGGGATAAAGGGCGAGATTTTGCCACAGTGCGCAAAGTATTGTCCATGCCCCAAG
TCAATATTGTCAAGAAAACAGAAGTACAGACAGGCGGATTCTCCAAGGAGTCAA
TTTTACCAAAAAGAAATTCGGACAAGCTTATTGCTCGTAAAAAAGACTGGGATCC
AAAAAAATATGGTGGTTTTGATAGTCCAACGGTAGCTTATTCAGTCCTAGTGGTT
GCTAAGGTGGAAAAAGGGAAATCGAAGAAGTTAAAATCCGTTAAAGAGTTACTA
GGGATCACAATTATGGAAAGAAGTTCCTTTGAAAAAAATCCGATTGACTTTTTAG
AAGCTAAAGGATATAAGGAAGTTAAAAAAGACTTAATCATTAAACTACCTAAAT
ATAGTCTTTTTGAGTTAGAAAACGGTCGTAAACGGATGCTGGCTAGTGCCGGAGA
ATTACAAAAAGGAAATGAGCTGGCTCTGCCAAGCAAATATGTGAATTTTTTATAT
TTAGCTAGTCATTATGAAAAGTTGAAGGGTAGTCCAGAAGATAACGAACAAAAA
CAATTGTTTGTGGAGCAGCATAAGCATTATTTAGATGAGATTATTGAGCAAATCA
GTGAATTTTCTAAGCGTGTTATTTTAGCAGATGCCAATTTAGATAAAGTTCTTAGT
GCATATAACAAACATAGAGACAAACCAATACGTGAACAAGCAGAAAATATTATT
CATTTATTTACGTTGACGAATCTTGGAGCTCCCGCTGCTTTTAAATATTTTGATAC
AACAATTGATCGTAAACGATATACGTCTACAAAAGAAGTTTTAGATGCCACTCTT
ATCCATCAATCCATCACTGGTCTTTATGAAACACGCATTGATTTGAGTCAGCTAG
GAGGTGACTAACTCGAGTAAGGATCTCCAGGCATCAAATAAAACGAAAGGCTCAG
TCGAAAGACTGGGCCTTTCGTTTTATCTGTTGTTTGTCGGTGAACGCTCTCTACTA
GAGTCACACTGGCTCACCTTCGGGTGGGCCTTTCTGCG

**RBP-AD gene:**
Activator proteins (SoxS-R93A/S101A) fused to RBP (MCP) are used in all the figures in the Main Manuscript and are expressed with the BBa_J23107 promoter from Table 2.

BBa_J23107, BBa_J34801, MCP-SoxS-R93A/S101A, BBa_B0015 terminator

T T T A C G G C T A G C T C A G C C C T A G G T A T T A T G C T A G C G A A T T C A T T A A A G A G G A G A A
A G G T A C C A T G G G G C C C G C T T C T A A C T T T A C T C A G T T C G T T C T C G T C G A C A A T G G C
G G A A C T G G C G A C G T G A C T G T C G C C C C A A G C A A C T T C G C T A A C G G G A T C G C T G A A T
G G A T C A G C T C T A A C T C G C G T T C A C A G G C T T A C A A A G T A A C C T G T A G C G T T C G T C A
G A G C T C T G C G C A G A A T C G C A A A T A C A C C A T C A A A G T C G A G G T G C C T A A A G G C G C
C T G G C G T T C G T A C T T A A A T A T G G A A C T A A C C A T T C C A A T T T T C G C C A C G A A T T C C
G A C T G C G A G C T T A T T G T T A A G G C A A T G C A A G G T C T C C T A A A A G A T G G A A A C C C G A
T T C C C T C A G C A A T C G C A G C A A A C T C C G G C A T C T A C G G T G G C G G A G G T A G C A T G T C
C C A T C A G A A A A T T A T T C A G G A T C T T A T C G C A T G G A T T G A C G A G C A T A T T G A C C A G
C C G C T T A A C A T T G A T G T A G T C G C A A A A A A A A T C A G G C T A T T C A A A G T G G T A C T T G C
A A C G A A T G T T C C G C A C G G T G A C G C A T C A G A C G C T T G G C G A T T A C A T T C G C C A A C G
C C G C C T G T T A C T G G C C G C C G T T G A G T T G C G C A C C A C C G A G C G T C C G A T T T T T G A T
A T C G C A A T G G A C C T G G G T T A T G T C T C G C A G C A G A C C T T C T C C C G C G T T T T C G C G C
G G C A G T T T G A T C G C A C T C C C A G C G A T T A T C G C C A C C G C C T G T A A G C G G C C G C C A C
G C A A A A A A C C C C G C T T C G G C G G G G T T T T T T C G C

##### 4.2 scRNA sequences

The base scRNA sequence used is 1XMS2 scRNA.b1, which has the dCas9 binding site and the MS2 RNA recruitment hairpins[5]. Here are the sequences of scRNAs used in various figures in the paper.

Target DNA binding site, dCas9 binding site, RBP-AD binding site, tracr RNA terminator

**scRNA**$_{\mathrm{RFP}}$ (J306 binding site):
TTGTGTCCAGAACGCTCCGTGTTTTAGAGCTAGAAATAGCAAGTTAAAATAAGGC
TAGTCCGTTATCAACTTGAAAAAGTGGCACATGAGGATCACCCATGTGCTTTTTTT

**scRNA**$_{\mathrm{GFP}}$ (J108 binding site, binding with dCas9 and RBP-AD):
TGGTAACCGCAGGACACCGCGTTTTAGAGCTAGAAATAGCAAGTTAAAATAAGGC
TAGTCCGTTATCAACTTGAAAAAGTGGCACATGAGGATCACCCATGTGCTTTTTTT

**Scrambled scRNA**$_{\mathrm{GFP}}$ (J108 binding site, no dCas9 or RBP-AD binding, used in Main Manuscript Fig. 3 and case (IV) in Main Manuscript Fig. 4):
TGGTAACCGCAGGACACCGCCATCTATGAGTTATGAGGTTAGGTTTTAGAGCTAG
GCAAATGTAAAACAAACATGTCCATCGGTTGCCCAAACACCCATGTGCTTTTTTT

**scRNA**$_{\mathrm{GFP}}$ (J108 binding site, binding with RBP-AD only, used as Case (II) in Main Manuscript Fig. 4):
TGGTAACCGCAGGACACCGCGTTTTAGAGCTAGAAATAGCAAGTTAAAATAAGG
CTAGTCCGTTATCAGCAGAGGCGATAAACATGAGGATCACCCATGTGCTTTTTTT

**scRNA**$_{\mathrm{GFP}}$ (J108 binding site, binding with dCas9 only, used as Case (III) in Main Manuscript Fig. 4):
TGGTAACCGCAGGACACCGCGTTTTAGAGCTAGAAATAGCAAGTTAAAATAAGGC
TAGTCCGTTATCAACTTGAAAAAGTGGCACCGAGTCGGTGCGCCATACTTTTTTT

**No scRNA**$_{\mathrm{GFP}}$ (promoter followed by terminator used in Main Manuscript Figs. 3 and 6):
pTrc promoter, L3S3P12 Terminator

#### References


1. A. J. Meyer, T. H. Segall-Shapiro, E. Glassey, J. Zhang, and C. A. Voigt, "Escherichia coli "Marionette" strains with 12 highly optimized small-molecule sensors," *Nature Chemical Biology*, vol. 15, no. 2, pp. 196–204, 2019.
2. M. G. Thompson, N. Sedaghatian, J. F. Barajas, M. Wehrs, C. B. Bailey, N. Kaplan, N. J. Hillson, A. Mukhopadhyay, and J. D. Keasling, "Isolation and characterization of novel mutations in the psc101 origin that increase copy number," *Scientific Reports*, vol. 8, no. 1, p. 1590, 2018.
3. K. Manoj and D. Del Vecchio, "Emergent interactions due to resource competition in CRISPR-mediated genetic activation circuits," in *2022 IEEE 61st Conference on Decision and Control (CDC)*, pp. 1300–1305, IEEE, 2022.
4. D. Del Vecchio and R. M. Murray, *Biomolecular feedback systems*. Princeton University Press Princeton, NJ, 2015.
5. J. Fontana, C. Dong, C. Kiattisewee, V. P. Chavali, B. I. Tickman, J. M. Carothers, and J. G. Zalatan, "Effective CRISPRa-mediated control of gene expression in bacteria must overcome strict target site requirements," *Nature Communications*, vol. 11, no. 1, p. 1618, 2020.
